# Molecular size dominates α_2_-adrenergic subtype-selectivity benchmarks: five controls for reducing attrition in selective ligand design

**DOI:** 10.64898/2026.08.18.745649

**Authors:** Manal A. Nael, Khaled M. Elokely

**Author notes:** Corresponding author: Khaled M. Elokely.

## Abstract

**Background:** Subtype-selectivity predictions are scored against measured selectivity and judged against an assumed noise ceiling. We asked what an α_2_-adrenergic benchmark rewards and which controls change its interpretation.

**Research design and methods:** On a frozen benchmark of 586 paired α_2A_/α_2C_ compounds we evaluated Glide SP docking, CNN rescoring, ligand-only fingerprint models, receptor descriptors and pose contacts, with dopamine D_3_/D_2_ as comparator, applying five controls: a measured ceiling, a cluster-identity null, a nonselective reference, a same-receptor floor and a trivial-descriptor baseline.

**Results:** Five descriptors from SMILES reached Spearman 0.645, 72% of the measured ceiling, against 0.071 for Glide SP and 0.188 for CNN rescoring; receptor properties and pose contacts reduced to size under control, while a non-size signal of 0.258 survived. Measured rather than propagated noise raised that ceiling from 0.704 to 0.897; cluster identity alone reached R^2^ 0.499 on D_3_/D_2_ and none on α_2_; a nonselective reference received +1.43 to +4.79 kcal/mol where zero is expected; and a same-receptor floor reached 1.77-fold against 1.88-fold across subtypes.

**Conclusions:** Such benchmarks reward molecular size first; a method must exceed 0.645 before its score indicates structural reasoning. The controls are inexpensive; conclusions rest on two receptor pairs, a three-pair floor and static structures.

## Introduction

Selectivity among closely related receptor subtypes decides whether a candidate advances or is set aside, and among the α_2_-adrenergic receptors that decision is currently taken with limited information. The three subtypes carry pharmacologically distinct actions: α_2A_ mediates the sympatholytic and withdrawal-suppressing effects that make clonidine and lofexidine useful in the management of opioid withdrawal^1^, while α_2B_ contributes the peripheral vasoconstrictive liability that constrains dosing. Separating the subtypes cleanly is the design objective, and it is the property a selectivity prediction is asked to supply.

The same receptor class has acquired a second and more urgent relevance. Xylazine, a veterinary α_2_ agonist, is now a widespread adulterant of the illicit fentanyl supply and is associated with a withdrawal syndrome and a wound pathology that opioid pharmacology alone does not explain^2^, and medetomidine has since appeared in the same supply^3^. Neither compound has a solved complex with any α_2_ subtype, and the attribution of their effects across subtypes remains an open pharmacological question. Any countermeasure begins with assigning activity to a subtype, and that assignment currently rests on prediction rather than on structures.

A computed selectivity score arrives long before any binding data, it is inexpensive, and it is often the only quantitative discriminator available for ordering a list of analogues. Two approaches carry most of the traffic: structure-based scoring, in which a ligand is docked into each subtype and the score difference is read as a preference, and ligand-based modeling, in which measured selectivity is regressed on chemical descriptors. Both are ordinarily evaluated by correlating predicted against measured selectivity, and both are ordinarily judged against an implicit ceiling, the level of performance that measurement error would permit. Synthesis decisions are taken on the strength of that comparison.

Whether such a score can bear that weight depends on five quantities that are usually assumed rather than measured. The first is the ceiling itself: selectivity is a difference of two affinities, and when both come from the same publication, as they usually do, their errors are correlated and partly cancel, so a propagated ceiling sits too low. The second is what a benchmark actually tests, since a model can score well by recognizing which chemical series a compound belongs to. The third is what a selectivity free-energy protocol returns for a ligand that has no selectivity, which bounds how much of any result is method rather than signal. The fourth is what a structural comparison returns for two structures of the same receptor, where subtype divergence is zero by construction. The fifth is what a handful of trivial descriptors achieve on the same benchmark, which sets the baseline a method must actually beat.

Here we assemble a single frozen, leakage-audited, provenance-tracked benchmark of 586 paired α_2A_/α_2C_ compounds, evaluate docking, CNN rescoring and ligand-only modeling against it, and measure each of those five quantities rather than assume it. The α_2A_ and α_2C_ crystal structures used throughout^4, 5^ were solved in the same state with the same antagonist bound, which makes the structural comparison unusually clean. Glide SP reaches a Spearman correlation of 0.071 with measured selectivity and CNN rescoring 0.188, while a ligand-only fingerprint model that is never shown a receptor reaches 0.564 to 0.741 depending on how the cross-validation is grouped. The structural reason is visible in the coordinates: of the 18 residues within 5.0 Å of the co-crystallized antagonist, 16 are shared between the subtypes, and the two that differ, Glu189→Gly203 and Ile190→Leu204, lie adjacent at the extracellular rim of the pocket. Five descriptors computable from a SMILES string reach Spearman 0.645, or 72% of the measured ceiling, so a method scored on a benchmark of this kind competes first of all against molecular size.

What a discovery team takes from this is a set of five controls, each inexpensive relative to the calculation it qualifies, and a specific target in place of an open one: a sufficient method must exceed 0.645 from trivial descriptors and must recover part of the 0.258 of non-size chemical signal that every structural readout tested here misses.

## Research design and methods

### Receptor structures and structural comparison

Crystal structures of the human α_2A_-(PDB 6KUX)^4^ and α_2C_-adrenergic receptor (PDB 6KUW)^5^ were used throughout. Both were solved in the inactive state bound to the same antagonist, RS-79948, deposited as ligand E3F in 6KUX and as its C13a epimer E33 in 6KUW. For every structural analysis we retained chain A only, discarded all heteroatoms and waters, and removed the crystallization fusion, which in both entries is numbered from residue 1001 upward. This left 269 modeled residues for α_2A_ and 278 for α_2C_. Author numbering is used in the text and is identical to UniProt numbering for both receptors; the DRY motif is D130-R131-Y132 in α_2A_ and D148-R149-Y150 in α_2C_, and Asp3.32 is Asp113 and Asp131 respectively, in the Ballesteros-Weinstein scheme^6^.

Superposition was performed in PyMOL^7^. We report both available estimates because they answer different questions: align, which builds a sequence alignment before fitting, converged after five cycles of outlier rejection to 0.638 Å over 1,425 atom pairs, from 2.093 Å over 1,907 pairs before rejection; super, which is sequence-independent and driven by structural superposition, gave 0.743 Å over 1,209 atom pairs. The sequence alignment underlying these fits contains 234 aligned residue pairs, of which 180 are identical (76.9%).

The orthosteric shell was defined without reference to any prior annotation, as the complete set of residues with at least one heavy atom within 5.0 Å of the co-crystallized RS-79948 in 6KUX. Whole residues were selected; taking atoms within the cutoff and then filtering to Cα would retain only those residues whose Cα itself falls inside the shell and would badly under-count it. This gives 18 residues. Divergence was assessed by mapping each shell residue through the sequence alignment to its α_2C_ counterpart.

To describe where divergent positions sit along the membrane normal, we projected the Cα coordinates of the transmembrane helices onto their first principal component. Helical secondary structure was assigned by PyMOL’s dss algorithm; loops and termini were excluded because they displace the principal axis away from the bundle axis. The sign was fixed by requiring the ECL2 centroid to lie above the DRY centroid. Each residue was then assigned a normalized axial coordinate

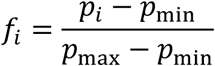

with p_i_ the projection of residue i, so that f = 0 is the intracellular and f = 1 the extracellular extreme. Figures were rendered from coordinates pre-rotated into this membrane frame, so that the deposited coordinate files are exactly those shown.

### Bioactivity data and the frozen evaluation set

Earlier analyses in this project were computed on three different compound sets drawn from two curations, and were therefore not comparable. The intersection of all three contained only 31 compounds, and on those 31 the two curations disagree on ΔpK_i_ by a mean of 0.267 log units (median 0.240, maximum 0.794); only 4 of the 31 agree to within 0.01 log units. Curation choice therefore contributes label disagreement of the same order as measurement error, and all results below are reported on a single frozen evaluation set.

The frozen set comprises 586 compounds with paired human α_2A_ and α_2C_ K_i_ values from ChEMBL^8^, built from 2,013 α_2A_ and 1,842 α_2C_ activities. Document-level provenance (document_chembl_id) was retained for every activity, so the noise ceiling, the cross-validation grouping and every method score are computed on the same compounds, the same labels and the same provenance. ChEMBL target identifiers were confirmed against UniProt accessions rather than assumed: CHEMBL1867 = α_2A_ (P08913), CHEMBL1942 = α_2B_ (P18089), CHEMBL1916 = α_2C_ (P18825).

A second, independent curation from BindingDB^9^ (108 compounds, of which 106 carry the quantities needed for the ceiling) was retained solely to test whether the ceiling correction replicates across databases. The dopamine comparison set comprises 934 paired human D_3_/D_2_ K_i_ compounds assembled under identical species, endpoint and unit filters (575 D_3_-preferring, 5 D_2_-preferring, 354 nonselective).

### The selectivity endpoint and its attainable ceiling

Subtype selectivity is a difference endpoint,

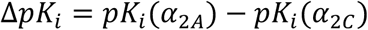

and its error structure is not that of a single affinity. Writing the observed values as pK_i_(α_2A_) = μ_A_ + ε_A_ and pK_i_(α_2C_) = μ_C_ + ε_C_, the variance of the difference is

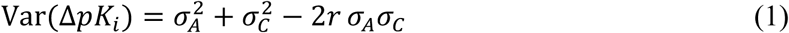

where r is the correlation between ε_A_ and ε_C_. The convention we set out to test is the one obtained by assuming r = 0 and σ_A_ = σ_C_ = σ, giving the propagated estimate

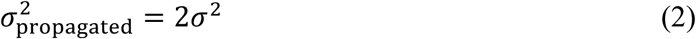

That assumption is not satisfied here. Paired affinities are predominantly co-reported: 525 of 586 compounds (90%) in the frozen set carry an α_2A_ and an α_2C_ value from the same source document, and so share laboratory, protocol, batch and radioligand. We therefore measured the noise directly rather than propagating it. For a compound measured in two independent documents i and j, the difference of its two reported selectivities, Δ_i_ − Δ_j_, contains only error, and

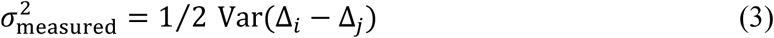

Equation (3) requires no assumption about r. Its unbiasedness was confirmed by Monte Carlo simulation (0.16532 recovered against a 0.16503 target over 4,000 replicates). In the frozen set, 55 compounds carry ≥ 2 paired documents and contribute 136 cross-document pair-differences, giving σ^2^_measured_ = 0.139 against σ^2^_propagated_ = 0.360 (σ = 0.424). In the BindingDB set the corresponding values are 0.165 against 0.366, from 24 compounds and 39 pair-differences. The within-document error correlation implied by these quantities is r = 0.659 and r = 0.633 respectively (Figure S2).

The attainable ceiling follows from classical attenuation^10^. With observed label variance σ^2^_obs_ and noise variance σ^2^_noise_, the maximum correlation any predictor can achieve against the noisy labels is

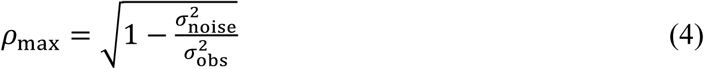

Substituting the measured and propagated noise variances gives ρ_max_ = 0.897 (bootstrap 95% CI 0.844 to 0.960) and 0.704 on the frozen set, and 0.819 (0.715 to 0.944) and 0.520 on the BindingDB set. Two caveats are carried explicitly. Equation (4) is a Pearson result while our reported scores are Spearman; simulation on the empirical ΔpK_i_ distribution shows the two differ by only 0.01 to 0.02 because the label distribution is close to normal, and the correction for median aggregation across multiple documents acts in the opposite direction and is slightly larger. And the confidence interval is computed over compound clusters (n = 24 clusters, ≈ 30 effective degrees of freedom on the BindingDB set), not over the individual (compound, document) deviations, which are not independent.

### Cross-validation, nulls and method scoring

All predictors were scored as Spearman rank correlations against measured ΔpK_i_ on the frozen set, with 95% confidence intervals from a compound-level cluster bootstrap.

The ligand-only null is a Morgan-fingerprint (ECFP-like, radius 2, 2,048 bits)^11^ random-forest^12^ model trained to predict ΔpK_i_ from ligand structure alone, with no receptor information of any kind. It is reported under three grouping schemes, because the grouping determines what the null actually measures: scaffold-grouped (ρ = 0.741, CI 0.698 to 0.784), document-grouped (0.599, 0.539 to 0.649), and grouped simultaneously on scaffold and document (0.564, 0.503 to 0.614). Document-grouped cross-validation is included specifically to exclude the possibility that a fingerprint model partially recovers document identity rather than chemistry.

The scaffold-grouped value was regenerated from the shipped frozen set with the forest fixed at 500 trees and seed 0, resampling compounds for the interval. Refitting across ten fold assignments gives 0.742 ± 0.002, so the estimate is stable to how the folds fall; resampling scaffold groups rather than compounds widens the interval to 0.655 to 0.809. The document-grouped schemes require source-document identifiers, which are not redistributable, and those two values are carried from the original curation.

### Docking

Receptors were prepared in Schrödinger 2025-2 (build 133; Glide 107133, mmshare 70133). Grids were centered on the co-crystallized ligand with a 12 Å inner box and a 30 Å outer box. α_2A_/α_2C_ docking of the frozen set used Glide standard precision (PRECISION SP)^13^. The dopamine D_3_/D_2_ calculation used Glide extra precision^14^.

CNN rescoring used GNINA 1.3.2^15^ over the full frozen set, 2,930 poses per receptor. We note that an earlier value of ρ = 0.27 for this method was obtained on the smaller 108-compound BindingDB-labeled subset; re-scored on the frozen set the value is 0.188 (CI 0.107 to 0.263), and the two intervals do not overlap, so only the frozen-set value is used.

Two competence controls were run before any scoring result was interpreted. In the first, five α_2_ co-crystal complexes were redocked blind, with the search box derived from the contact frame rather than from the native ligand: 7W7E (α_2A_, active, biased agonist), 9CBL (α_2A_, active, epinephrine), 6KUX (α_2A_, inactive, RS-79948), 6K41 (α_2B_, active, dexmedetomidine) and 6KUW (α_2C_, inactive). Median top-ranked symmetry-corrected heavy-atom RMSD values were 0.37, 0.25, 0.63, 0.42 and 0.73 Å, with all 20 of 20 seeds within 2 Å for every complex. In the second, conformational state was assigned from a prespecified structure-only coordinate, the TM3-TM6 distance measured between the Cα atoms of positions 3.50 and 6.34, against a threshold of 10 Å fixed in advance from the literature: active structures gave 15.92 to 16.08 Å and inactive structures 6.31 to 6.83 Å, with all three blinded structures called correctly.

Both controls establish search and setup competence only. Self-redocking does not measure prospective accuracy: the docking engine may have been exposed to these structures during development, and the search box is defined by contacts of the native ligand. They are reported to exclude setup as an explanation for the scoring results, not as evidence of predictive performance.

### Chemotype separability

To quantify how much of a selectivity benchmark is solvable without any structural or chemical reasoning, compounds were grouped by Butina clustering^16^ of Morgan fingerprints at a Tanimoto cutoff of 0.35, applied identically to both receptor pairs. Three measurements were made.

T1, the decisive one, is a model that receives only a one-hot encoding of cluster membership (no chemistry, no structure, nothing but which series a compound belongs to) and is scored out-of-fold. T_2_ compares an ECFP model under random-split against cluster-grouped cross-validation, the difference indicating within-series memorization. T_3_ is a cluster-identity-only classifier of the selective class, scored by AUC.

The α_2_ set contains 108 compounds in 89 clusters (1.2 compounds per cluster) and the D_3_/D_2_ set 934 in 226 clusters (4.1 per cluster). Because the α_2_ set is close to one compound per cluster, the between-cluster variance fraction η^2^ is near-total by construction (0.936 against 0.773) and is therefore reported only for completeness and not used as a test; T_1_ and T_3_ are cross-validated and immune to that artifact.

### Contact divergence and the same-receptor floor

Contact networks^17^ were computed for eight receptor pairs under a single harmonized construct definition: chain A only, heteroatoms removed and crystallization fusions at residue ≥ 1000 stripped. The extracellular region was defined geometrically and identically for every pair, as the top third of the helical-bundle principal axis, with orientation fixed by the N-versus C-terminal position; no per-system tuning was applied. Enrichment is

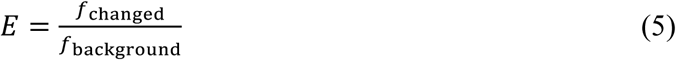

the fraction of subtype-divergent contacts falling in the extracellular region divided by the same fraction over the receptor-wide background contact set. Significance was assessed by a permutation null that resamples the observed background contact set to the size of the changed-contact set (20,000 draws).

The geometric rule was validated in two ways before being applied. It reproduces a hand-defined analysis of the α_2_ pair (1.88 against 1.79; p = 0.00005 against 0.00015), so it is not tuned to produce the α_2_ answer; and applied unchanged to D_3_ versus D_2_, where a secondary binding pocket is independently established as the selectivity determinant, it localizes divergence to the extracellular region at 1.60-fold.

The panel comprises five cross-subtype pairs and three same-receptor pairs. The same-receptor comparisons, built from two independent crystal structures of one receptor, supply a technical floor whose true subtype divergence is zero by construction.

### Target-side structural profiling and its floor

Both receptors were profiled as apo structures with a contact-network engine, using the same harmonization as every other structural analysis here: chain A only, heteroatoms and waters removed, crystallization fusions at residue >= 1000 stripped. This gives global descriptors (buried void volume, cavity count, maximum travel depth, total contact energy, total sidechain entropy, surface electrostatic statistics), per-pocket geometry, and per-residue channels for dynamic coupling, contact energy, sidechain entropy, travel depth, coordination, rigidity and entanglement.

Because any structural quantity measured on one pair of structures is subject to the objection raised in Calibration 4, the identical profiling was applied to three pairs of independent structures of a single receptor, where the true difference is zero by construction: β_2_ (2RH1 and 3NY8), 5-HT_2B_ (4IB4 and 4NC3) and A_2A_ (3EML and 4EIY). Quantities that scale with chain length, total contact energy and total sidechain entropy, are reported per residue so that pairs of differing modeled length remain comparable.

A first profiling run was discarded before analysis. It used the deposited entries without harmonization, and because 6KUW contains two receptor copies and both entries retain their fusions, α_2C_ returned 946 residues against α_2A_’s 375; every global comparison from that run described the contents of the crystallographic file rather than the receptor.

### Joining receptor properties to docked poses

Per-compound descriptors were formed by combining the per-residue receptor properties with each ligand’s own contact profile. For every compound the best-scoring Glide pose in each receptor was taken, verified by reproducing the archived docking scores exactly for all 586 compounds, and the number of ligand heavy atoms within 4.5 Å of each receptor residue recorded. Contacts were computed in the coordinate frame Glide docked into; the copy of α_2C_ used for profiling and for figures had been superposed onto α_2A_ and sits 82.7 Å away, so the two were joined by residue number, which is frame-independent, rather than by coordinate.

Two constructions were tested. The first sums each property over the contacted residues and differences the two receptors, which admits ligand size; the second records the per-residue contact pattern without summing. Both were scored by Spearman correlation against measured dpKi under scaffold-grouped cross-validation, against three controls: a size-only baseline of six contact-count features, a permutation that reassigns property values across residues while leaving contacts untouched, and a column shuffle that permutes residue identity within each compound and so preserves that compound’s total contact count exactly.

### Removing molecular size

Size was represented three ways of increasing strength: molecular weight alone; molecular weight with heavy-atom count; and a block of five descriptors adding rotatable bonds, topological polar surface area and calculated logP. Residualization was performed inside the cross-validation loop, fitting the size model on the training fold only and predicting the residual on both folds, so that no information from the held-out fold reaches the feature model through the size model. Residualizing on the full set before splitting would leak exactly the quantity under test.

### Free-energy calculation

Potentials of mean force along a one-dimensional ligand-receptor distance coordinate were computed for D_3_ and D_2_ by well-tempered metadynamics^18^ with three walkers per system^19^, 150 ns per walker (450 ns per system). Free energies were extracted across nine analysis definitions, formed by crossing three bulk reference distances (1.8, 1.9, 2.0 nm) with three orthosteric boundaries (0.8, 0.9, 1.0 nm), and the binding free energy taken as the well depth. The nonselective reference ligand was eticlopride and the selective comparator SB-277011A (PubChem CID 5311096)^20^.

This calculation is reported as a design requirement rather than as a measurement, and five specific defects are documented in the Results. Of these, two are relevant to reproducing the setup: the three walkers shared a single bias grid, so their mutual agreement is near-perfect by construction and is not a convergence diagnostic; and the free-energy surface was overwritten every 100 ps, so only the final surface was retained and no time-block convergence test is possible from the saved artifacts.

### Software

Structure handling and rendering used PyMOL. Cheminformatics and machine learning used Python 3.11.13 with RDKit 2025.09.5, scikit-learn 1.8.0, NumPy 2.4.2 and pandas 3.0.1. Docking used Schrödinger and GNINA 1.3.2. Figures were produced with Matplotlib 3.11.1^21^. Cheminformatics used RDKit^22^. All calculations were run on the University of Wyoming Advanced Research Computing Center MedicineBow cluster.

## Results

### The two receptors differ at two positions in the ligand shell, and both sit at its extracellular rim

α_2A_ and α_2C_ are close enough in the inactive state that the comparison is almost uninformative at the level of the fold. Superposed on structure alone, they agree to 0.743 Å over 1,209 atom pairs; with a sequence alignment imposed first and outliers rejected, to 0.638 Å over 1,425 (Figure S1). Of 234 aligned residue pairs, 180 are identical.

The interesting comparison is local. Defining the orthosteric shell without reference to any prior annotation, as every residue with a heavy atom within 5.0 Å of the co-crystallized antagonist RS-79948, gives 18 residues (Figure 1a). Sixteen of these are shared between the subtypes. The aminergic anchor is among the conserved: Asp3.32 is present in both, positioned identically, and every residue that contacts the ligand’s cationic center is common to the two receptors. If subtype selectivity were written into the walls of the orthosteric pocket, there is very little here to write it with.

**Figure 1.**
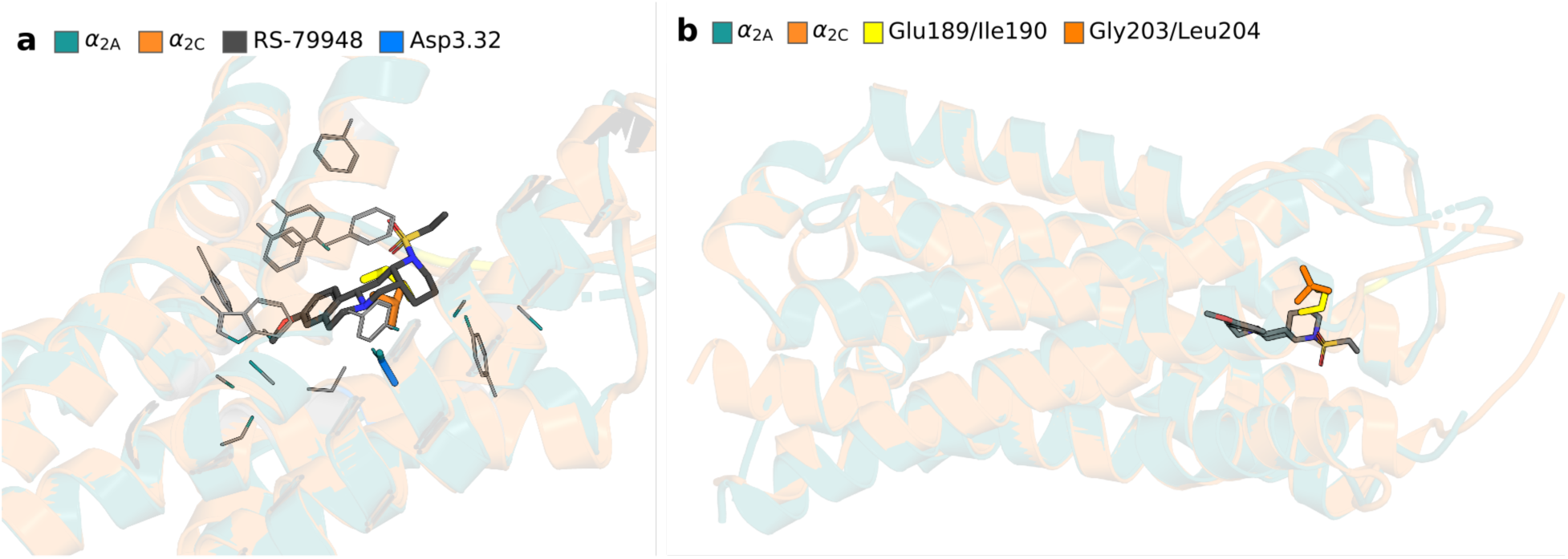
The orthosteric shell of α_2A_ and α_2C_ is almost entirely conserved, and what differs sits at its extracellular rim. (a) The 18 residues with at least one heavy atom within 5.0 Å of the co-crystallized antagonist RS-79948 in α_2A_ (6KUX). The ligand is shown as dark sticks and the shell residues as thin grey lines; the conserved Asp3.32 anchor is blue. The two positions that differ between the subtypes are shown as thick sticks, α_2A_ in yellow (Glu189, Ile190) and α_2C_ in orange (Gly203, Leu204). Cartoons are α_2A_ in teal and α_2C_ in orange, both at 85% transparency. (b) The same superposition viewed from the extracellular side, with the divergent positions in the same colors. Every canonical aminergic contact (Asp3.32, both TM5 serines, the Trp6.48 toggle, the Phe6.51/Phe6.52 aromatic cage, Tyr6.55 and Tyr7.43) is shared between the two receptors; the complete residue-by-residue comparison is given in Table S1.

The two positions that do differ are Glu189→Gly203 and Ile190→Leu204, and they are adjacent, at the top of the pocket where it opens into the extracellular vestibule (Figure 1a, yellow and orange). This is not a general property of the divergent residues: it is where they concentrate. Projecting every residue onto the transmembrane bundle axis and normalizing so that 0 is the intracellular and 1 the extracellular extreme, divergent positions have a mean axial coordinate of 0.528 against 0.478 for the receptor as a whole, and 38.9% of them fall in the extracellular third against 27.5% of all residues (Figure 1b). The divergence is modestly but consistently displaced toward the extracellular face, and inside the ligand shell it is confined entirely to the rim.

This sets up the question the rest of the paper addresses. The subtypes differ where a ligand would first encounter them and barely at all where it finally binds, so any method that scores a static pose in the orthosteric pocket is being asked to detect a difference that is largely not there.

### Static structure-based scoring does not rank α_2_ subtype selectivity

On the frozen set of 586 paired compounds, Glide SP reaches a Spearman correlation with measured ΔpK_i_ of 0.071 (95% CI −0.007 to 0.151); the interval includes zero. CNN rescoring with GNINA improves this to 0.188 (0.107 to 0.263) but no further (Figure 2).

**Figure 2.**
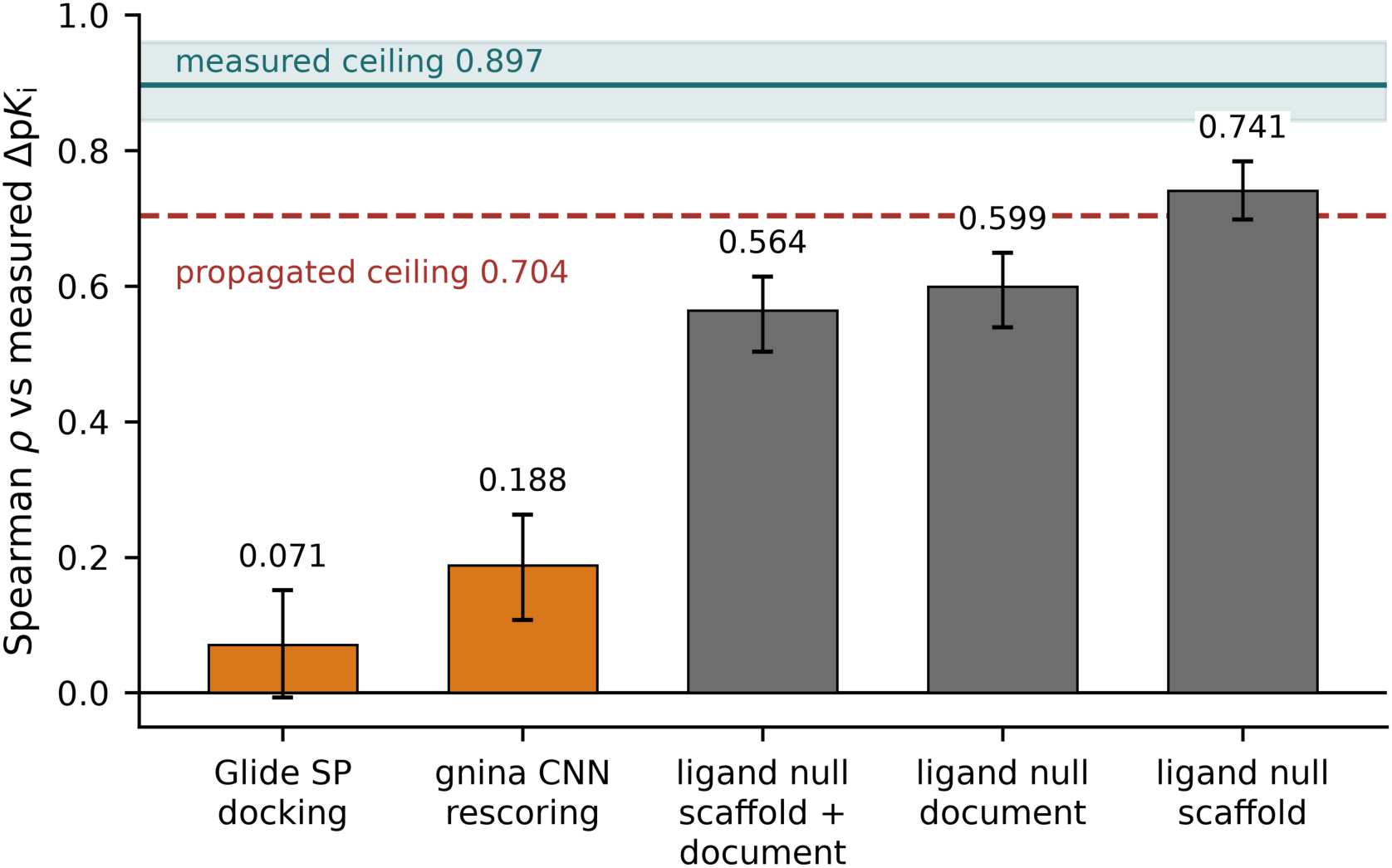
On a single frozen evaluation set, both docking methods are outperformed by a model that never sees a receptor. Spearman rank correlation against measured ΔpK_i_ for 586 paired α_2A_/α_2C_ compounds. Orange, structure-based methods; grey, the ligand-only fingerprint null under three cross-validation groupings. Error bars are 95% confidence intervals from a compound-level cluster bootstrap. The teal line and band mark the ceiling measured directly from replicate data (0.897; 95% CI 0.844 to 0.960); the red dashed line marks the ceiling obtained by propagating single-endpoint assay noise under an independence assumption (0.704).

The comparison that matters is not against zero but against a model that is given no structural information at all. A Morgan-fingerprint random forest, trained to predict selectivity from the ligand alone and never shown a receptor, reaches 0.741 under scaffold-grouped cross-validation. That number is inflated by the ease of the split, and under the two stricter groupings it falls to 0.599 when compounds sharing a source document are held out together, and 0.564 when scaffold and document are grouped simultaneously. Even the most conservative of these, 0.564, is three times the CNN score and eight times the Glide score. Both docking methods are beaten, comfortably, by a model that does not know the receptors exist.

Leakage-audited ligand QSAR bounds what chemistry alone supplies on the same receptor pair: R^2^ = 0.419 for α_2C_ versus α_2A_, with 0.370 for α_2A_/α_2B_ and 0.286 for α_2C_/α_2B_. Chemistry accounts for a substantial part of the signal and structure, as scored here, adds nothing on top of it.

### Setup is not the explanation

Before drawing any conclusion from the scoring results above, we established that the docking setup can do the things it would need to do. Five α_2_ co-crystal complexes were redocked blind, with the search box taken from the contact frame rather than from the native ligand. Median top-ranked RMSD values were 0.37 Å (7W7E, α_2A_ active, biased agonist), 0.25 Å (9CBL, α_2A_ active, epinephrine), 0.63 Å (6KUX, α_2A_ inactive, RS-79948), 0.42 Å (6K41, α_2B_ active, dexmedetomidine) and 0.73 Å (6KUW, α_2C_ inactive). All 20 of 20 seeds landed within 2 Å for every complex, across agonists and antagonists and all three subtypes.

A prespecified structure-only classifier, the TM3-TM6 Cα distance between positions 3.50 and 6.34 against a 10 Å threshold fixed in advance, separated active from inactive structures with a wide margin: 15.92 to 16.08 Å against 6.31 to 6.83 Å, calling all three blinded structures correctly.

Both are competence checks and we treat them as nothing more. Self-redocking measures search, not prospective accuracy; the engine may have been exposed to these structures, and the box is defined by the native ligand’s own contacts. What they establish is narrow and sufficient for our purpose: pose recovery and state assignment are not the reason the selectivity scores are low.

### Where the determinant is known, a static score still recovers only part of the magnitude

The α_2_ result could be dismissed as a case where nobody knows what the determinant is. Dopamine D_3_ versus D_2_ is the opposite case. The D_3_ secondary binding pocket is the textbook example of a structurally characterized subtype-selectivity determinant, and R-22 is the ligand whose co-crystal defined it^23^, at 566-fold D_3_ selectivity, or 3.906 kcal/mol at 310 K.

Glide XP scores R-22 at −9.580 kcal/mol against D_3_ and −7.971 against D_2_^24^, a predicted difference of −1.609 kcal/mol. Because a docking score is not a binding free energy, the more informative quantity is the same difference computed for a nonselective ligand and subtracted: eticlopride gives −6.969 and −6.551, a difference of −0.418. The baseline-corrected prediction is therefore −1.190 kcal/mol against a measured −3.906.

We deliberately do not convert this to a percentage. Treating a GlideScore difference as a free energy is the assumption under scrutiny, and expressing the shortfall as a fraction would import it. The statement the data supports is simpler: where the determinant is known, the pocket is characterized and the ligand is the one that defined it, a static score reproduces the direction and roughly a third of the magnitude.

### Calibration 1. The attainable ceiling must be measured, not propagated

A weak correlation is only interpretable against a ceiling, and for selectivity that ceiling is routinely obtained by propagating single-endpoint assay noise through the difference. That propagation assumes the two affinities carry independent errors. For this benchmark they do not: 525 of 586 compounds (90%) carry both of their values from the same source document, sharing laboratory, protocol, batch and radioligand.

Measuring the noise instead of assuming it changes the number substantially. Same-compound cross-document replicates give a selectivity noise variance of 0.139 log^2^ units on the frozen set, against 0.360 by propagation; the measured noise is 39% of the propagated value. The implied within-document error correlation is r = 0.659 (Figure S2). Through the attenuation relation this raises the attainable Spearman ceiling from 0.704 to 0.897 (bootstrap 95% CI 0.844 to 0.960) (Figure 3).

**Figure 3.**
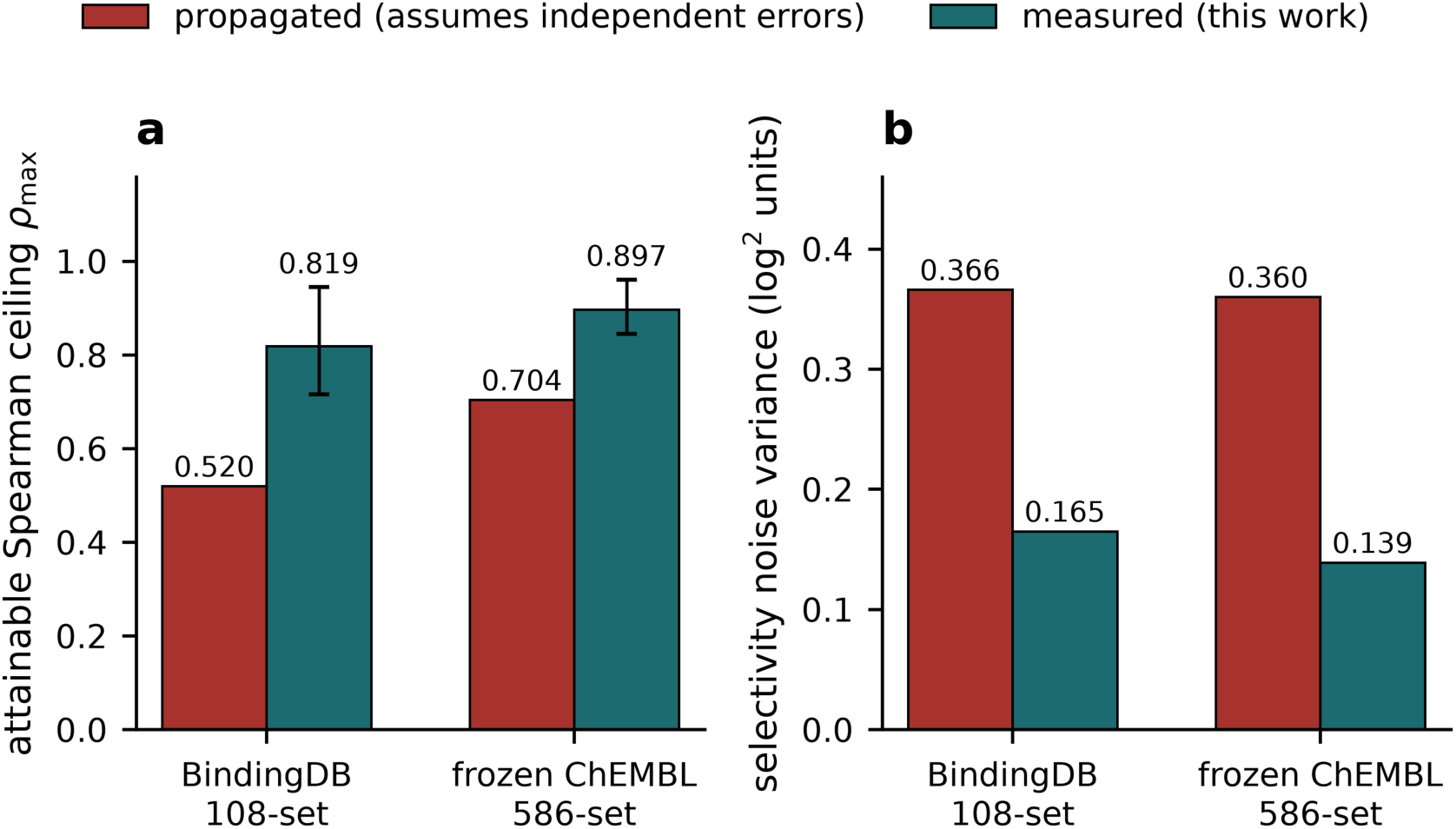
Measuring rather than propagating the noise raises the attainable ceiling, and the correction replicates across two independently assembled curations. (a) Attainable Spearman ceiling under the two estimators. Error bars are bootstrap 95% confidence intervals over compound clusters. (b) Selectivity noise variance under the two estimators. Red, propagated as 2σ^2^ assuming independent errors; teal, measured directly from same-compound cross-document replicates, which requires no assumption about the error correlation. The measured noise is 45% of the propagated value on the BindingDB set and 39% on the frozen ChEMBL set. The underlying within-document error correlation is shown in Figure S2.

The correction replicates on an independent curation. On the BindingDB set, assembled separately and with 2.3-fold less evidence, the measured noise variance is 0.165 against 0.366 propagated, r = 0.633, and the ceiling moves from 0.520 to 0.819 (0.715 to 0.944). The agreement between the two is a replication across databases, not an internal consistency check.

The consequence is that every gap in Figure 2 widens. Against the propagated ceiling, Glide SP recovers 10% of what is attainable and the ligand-only null appears to saturate it. Against the measured ceiling, Glide SP recovers 7.9%, GNINA 20.9%, and the strictest ligand-only null 62.9%, so the null does not saturate the ceiling either, and there is real headroom that neither chemistry nor static structure is reaching. An under-measured ceiling flatters weak predictors and conceals that headroom.

Two limits are worth stating. The attenuation relation is a Pearson result while the reported scores are Spearman; simulation on the empirical label distribution puts that discrepancy at 0.01 to 0.02, with the median-aggregation correction acting in the opposite direction and slightly larger. And the confidence interval is computed over compound clusters rather than over the individual (compound, document) deviations, which are not independent.

### Calibration 2. Chemotype separability, or why the standard benchmark misleads

D_3_/D_2_ is the convenient benchmark for selectivity prediction and ligand QSAR performs well on it. We asked whether that performance requires any chemical or structural reasoning at all, by training a model on nothing but cluster membership, a one-hot encoding of which Butina series a compound belongs to, containing no chemistry and no structure.

Series identity alone explains half the variance in D_3_/D_2_ selectivity (R^2^ = +0.499) and none of the variance in α_2_ selectivity (−0.017). The classifier tells the same story: cluster identity calls the D_3_-selective class at AUC 0.852 and the α_2_ class at 0.489, which is chance (Figure 4).

**Figure 4.**
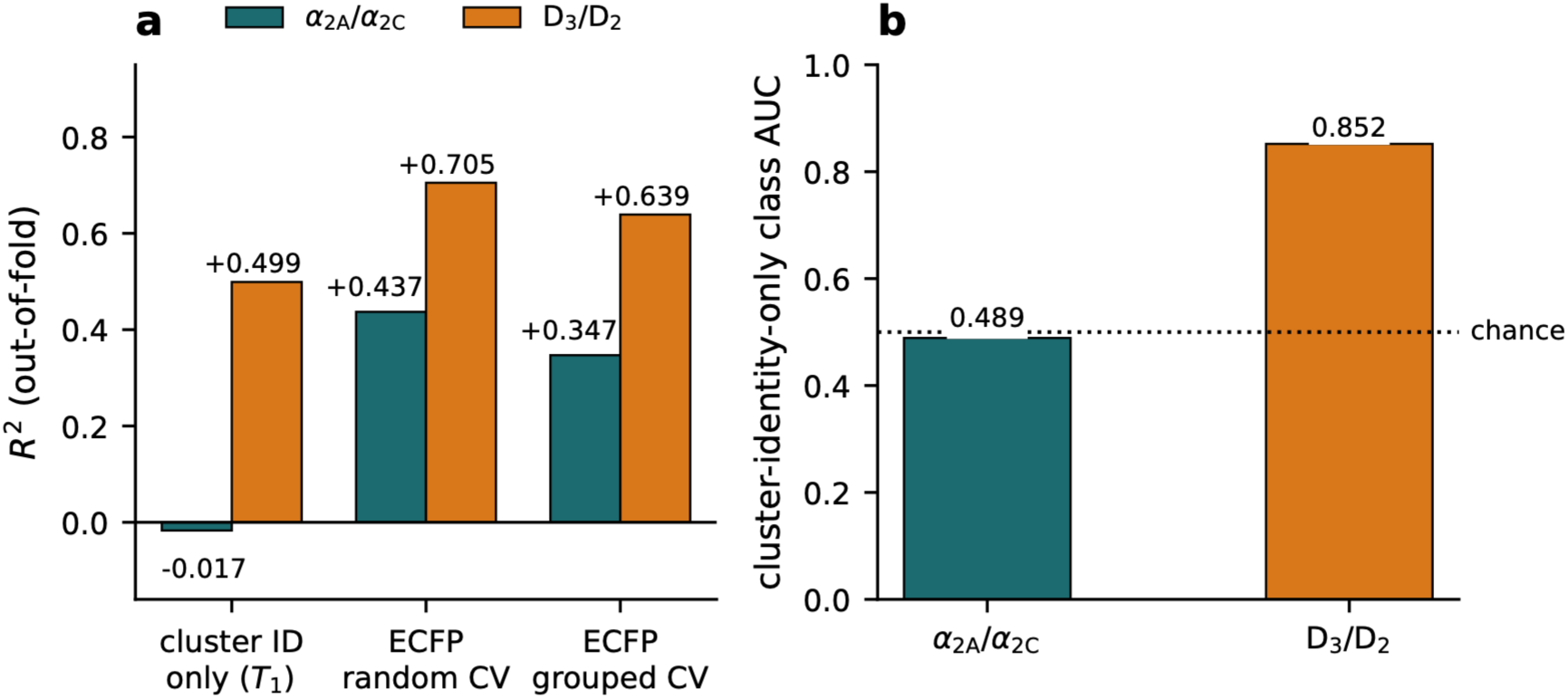
The dopamine benchmark is substantially solvable by series recognition and the α_2_ benchmark is not. (a) Out-of-fold R^2^ for three models. Cluster identity only (T1) is a model given nothing but a one-hot encoding of Butina cluster membership, with no chemistry and no structure. (b) Area under the ROC curve for a cluster-identity-only classifier of the selective class; the dotted line marks chance. Teal, α_2A_/α_2C_; orange, D_3_/D_2_. Full statistics, including the within-series memorization test and the between-cluster variance fraction, are given in Table S4.

The standard benchmark is therefore substantially solvable by series recognition, and a structure-based method evaluated on it is being scored on a task it does not need to solve. α_2_ is the regime where that shortcut is unavailable, which is why α_2_ is the informative case.

The within-series memorization test does not discriminate between the two pairs (the drop from random to cluster-grouped cross-validation is 0.090 for α_2_ and 0.066 for D_3_/D_2_) and is reported as a null. Between-cluster variance is uninformative at α_2_’s granularity, where 108 compounds occupy 89 clusters, and is not used as a test.

### Calibration 3. A selectivity protocol must return zero on a nonselective reference

A selectivity protocol is interpretable only if it returns zero for a ligand that has none. Running that reference through a one-dimensional distance-coordinate metadynamics PMF on D_3_/D_2_, and re-analysing across nine basin definitions, the nonselective control eticlopride receives between +1.43 and +4.79 kcal/mol of apparent selectivity where the true value is approximately zero. It is never near zero under any definition. SB-277011A, a genuinely D_3_-selective antagonist, receives −0.60 to +0.17, approximately zero, so the protocol inverts the ordering rather than merely adding noise.

The direction is robust and the magnitude is not. The 3.36 kcal/mol spread across analysis definitions is as large as the roughly 3.8 kcal/mol effect being modeled, so no single value from this run is quotable.

The defects are ours, and there are five. The free-energy surface was still filling at 150 ns per walker, and asymmetrically: over the final fifth of the run the span grew by about +1.6 kcal/mol in the D_3_ eticlopride system against +0.91 in D_2_, leaving roughly 0.7 kcal/mol of differential still accumulating when the run stopped. Only the final surface was retained, overwritten every 100 ps, so no time-block convergence test is possible. The walker dispersion one would normally quote, 0.004 to 0.058 kcal/mol, is not a convergence diagnostic here because the three walkers shared a single bias grid and their agreement is near-perfect by construction. The three D_3_ topologies carry no disulfide bonds at all: two cysteine pairs sit at 2.03 Å in the coordinates, Cys72-Cys150 (the conserved Cys3.25-CysECL2 bond) and Cys227-Cys230, yet neither is bonded in the force field, leaving 13 free thiols per system, and that bond tethers ECL2 to TM3 at the extracellular vestibule the reaction coordinate crosses. Finally, SB-277011A had no crystal pose; its geometry is a docked conformer, so its near-zero value cannot carry the inversion claim on its own.

We therefore claim nothing here about metadynamics, enhanced sampling or this collective variable as methods. These are five defects of one run, two of them bookkeeping errors that a topology check now catches. What survives is the design requirement, which is independent of all five: run a nonselective reference, and report the spread across analysis definitions, before quoting any selectivity free energy. Without the reference, an effect-sized number would have read as a success.

### Calibration 4. A structural-divergence statistic must clear a same-receptor floor

Contact-difference analyses between two receptor structures are used to localize selectivity determinants, and are conventionally tested against a permutation null that randomizes contact placement. Against that null the α_2_ result is emphatic: subtype-divergent contacts are enriched 1.88-fold in the extracellular region at p = 0.00005. Applied unchanged to D_3_ versus D_2_, where the answer is independently known, the same detector localizes divergence at 1.60-fold, so it is not simply manufacturing a signal.

The control that was missing is the same analysis applied to two independent structures of the *same* receptor, where subtype divergence is zero by construction. That floor is not 1.0. A_2A_ against A_2A_ gives 1.77-fold at p = 0.00055 and 5-HT_2B_ against 5-HT_2B_ gives 1.68-fold at p = 0.011 (Figure S3).

Placed against that floor, the α_2_ value of 1.88 is no longer distinguishable. The four interpreted cross-subtype pairs average 1.51 and the three same-receptor pairs average 1.54, and one genuine cross-subtype pair, M_2_ against M_3_, shows no enrichment at all (1.06, p = 0.36). Extracellular loops are the most conformationally variable region of any two class A GPCR structures, so changed contacts concentrate there whether or not the structures differ in subtype.

We therefore do not claim that this statistic demonstrates subtype-specific extracellular localization. A p-value against an inappropriate null is not evidence, and the appropriate null here is a technical floor rather than random placement. The β_1_/β_2_ pair is excluded from interpretation because β_1_ is the turkey receptor and its ECL2 landmark sits mid-axis, making its region assignment unreliable; three floor pairs is a small sample, and the conclusion rests on the existence of same-receptor values at 1.68 and 1.77, not on the floor mean.

This does not overturn the structural observations of the first section, which are independent measurements: the shell is 18 residues with 2 divergent, and the divergent positions are displaced extracellularly. What it removes is the claim that a contact statistic can establish that this displacement is subtype-specific rather than crystallographic.

### The receptors differ in interior packing, by margins three same-receptor pairs do not reproduce

Calibration 4 disqualified a contact statistic on a single structure pair. The same objection applies to any structural quantity measured on one pair, including the global descriptors used below, so the same control was built for them.

Both receptors were profiled independently as apo structures, chain A with crystallization fusions removed, giving buried void volume, cavity count, maximum travel depth, per-residue contact energy and sidechain entropy, and surface electrostatic statistics. The identical profiling was applied to three pairs of independent structures of a single receptor, where the true difference is zero by construction: β_2_ (2RH1, 3NY8), 5-HT_2B_ (4IB4, 4NC3) and A_2A_ (3EML, 4EIY).

| property | $\alpha_{2A}$ vs $\alpha_{2C}$ | $\beta_2$ | 5-HT<sub>2B</sub> | A<sub>2A</sub> |
| --- | --- | --- | --- | --- |
| buried void (Å<sup>3</sup>) | **166** | 33 | 25 | 87 |
| cavities $\geq 10$ Å<sup>3</sup> | **3** | 0 | 1 | 2 |
| maximum travel depth (Å) | **7.5** | 4.5 | 3.0 | 1.5 |
| surface potential standard deviation | **134.6** | 5.5 | 2.4 | 61.4 |
| contact energy per residue | **0.34** | 0.14 | 0.04 | 0.17 |
| sidechain entropy per residue | **0.08** | 0.01 | 0.00 | 0.03 |
| positive patch fraction | 0.01 | 0.10 | 0.24 | 0.01 |

Six of the seven cross-subtype differences exceed the largest same-receptor difference, by factors of 1.5 to 2.7 (Figure S4a). In absolute terms α_2C_ is the more cavernous receptor, with 355 Å^3 of buried void against 189, four cavities against one, and 7.5 Å more maximum travel depth, while α_2A_ carries the more electrostatically heterogeneous surface. Resolved position by position along the alignment, the difference in sidechain entropy is distributed across the sequence rather than concentrated at the two divergent orthosteric-shell positions (Figure S4b).

#### Three limits are carried with this result and none of them is cosmetic

The floor is three pairs, so exceeding its maximum has roughly a one-in-four chance per property under the null and about 1.8 of seven would be expected to clear by chance alone. The seven properties are not independent: buried void, cavity count and travel depth all report interior packing, so the effective number of tests is nearer two or three than seven. And the A_2A_ floor pair is itself heterogeneous, 288 modeled residues against 299, which inflates that floor and is why it is the largest in most rows.

The one property that does not clear is the informative one. Positive patch fraction differs by 0.01 between the subtypes while same-receptor pairs differ by up to 0.24, which shows the floor is not trivially small and that the exceedances are not an artifact of an over-tight control.

We therefore report this as an observation with a stated floor rather than as a demonstrated subtype-specific difference: the two receptors differ in interior packing by margins that three independent same-receptor comparisons do not reproduce, on a sample of three. It is offered as a target-side description, and no predictive claim rests on it. Nothing in the preceding sections depends on it either: the descriptors built from these properties, joined to the docked poses, reduce to molecular size under control.

### Calibration 5. Most of what the benchmark measures is molecular size

The four controls above ask whether a method’s score is interpretable. A fifth asks something simpler: what does the benchmark reward?

Molecular weight alone, with no model and no fitting, correlates with measured ΔpK_i_ at Spearman −0.561. Heavy-atom count gives −0.575. A block of five descriptors available from a SMILES string in milliseconds, molecular weight, heavy-atom count, rotatable bonds, topological polar surface area and calculated logP, reaches ρ = 0.645 under scaffold-grouped cross-validation, or 72% of the measured ceiling (Figure 5). Larger ligands are systematically less α_2A_-selective across this set.

**Figure 5.**
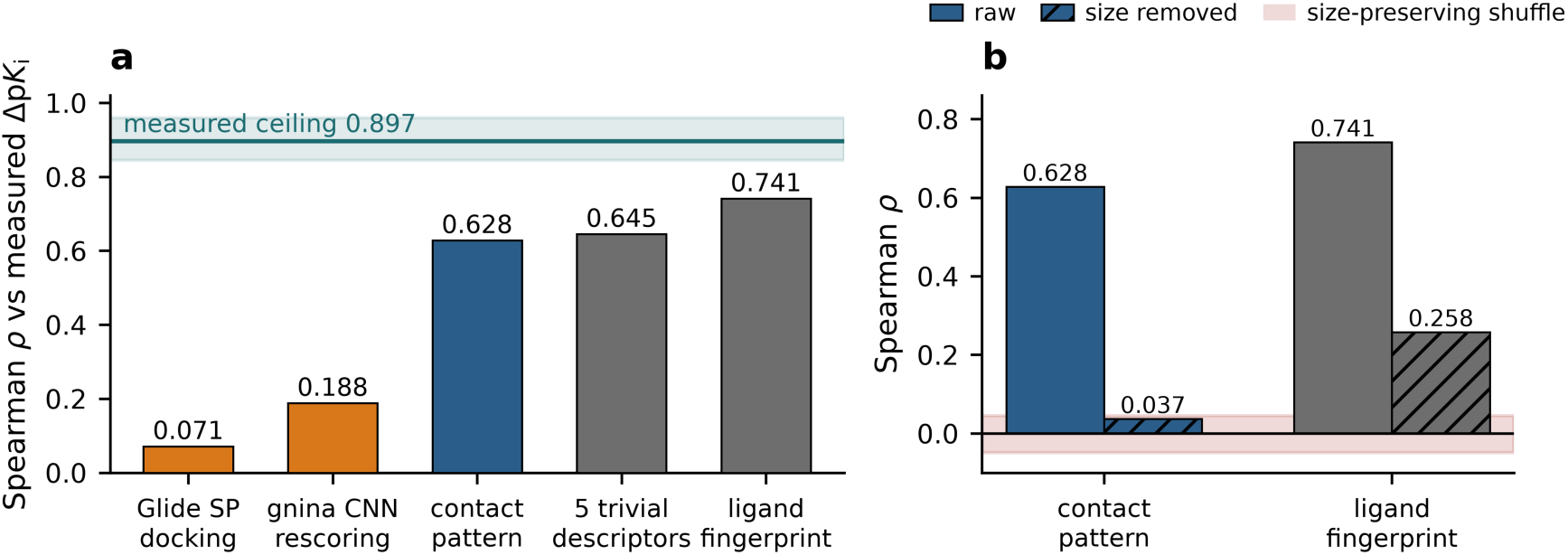
Molecular size accounts for most of what the benchmark measures, and structural features add nothing beyond it. (a) Spearman correlation with measured ΔpK_i_ on the frozen set. Orange, structure-based scoring; blue, the per-residue contact pattern taken from the same poses; grey, ligand-only baselines. The five trivial descriptors are molecular weight, heavy-atom count, rotatable bonds, topological polar surface area and calculated logP. Teal line and band, the measured ceiling. (b) The same two feature sets before and after molecular size is removed, with the size model fitted within each training fold and the residual predicted. Red band, the performance of a size-preserving shuffle of the contact pattern on the same residual (−0.002 ± 0.047). The contact pattern falls to 0.037 and enters that band; the fingerprint retains 0.258.

That single number reorders everything reported above. Five trivial descriptors outperform CNN rescoring (0.188) by more than threefold and Glide SP (0.071) by ninefold, and reach most of the way to the ligand-only fingerprint null (0.741). Any method scored on this benchmark is competing, first of all, against molecular size.

The consequence for interpretation is not that the benchmark is worthless but that its baseline is much higher than a correlation against zero implies. A structure-based method reporting ρ = 0.3 on a selectivity benchmark of this kind has not demonstrated structural reasoning; it has demonstrated less than molecular weight.

#### A non-size signal does exist, and ligand descriptors capture it

Removing size properly, by fitting the size model within each training fold and predicting the residual, leaves the Morgan fingerprint at ρ = 0.258, or 29% of the ceiling. Chemistry therefore knows something about α_2_ selectivity beyond how large a molecule is. The question the remaining sections answer is whether anything derived from receptor structure knows it too.

### Receptor properties and pose contacts add nothing beyond size

Docking scores discard most of what a pose contains, so we asked whether the poses themselves carry subtype information that the scoring function throws away. Both receptors were profiled independently with a contact-network engine, giving per-residue dynamic coupling, contact energy, sidechain entropy, travel depth, coordination, rigidity and entanglement, and these were joined to the docked poses of all 586 compounds through each ligand’s own contact profile. Every pose used is the one that produced the docking scores reported above, verified by reproducing the archived scores exactly.

Two feature constructions were tested, each against a control designed to break it.

#### Contact-weighted receptor properties

Summing each receptor property over the residues a ligand contacts, and differencing between subtypes, gives ρ = 0.406, which is 5.7-fold the Glide SP score computed from the identical poses. The result does not clear its control. Six features that count contacts and nothing else reach 0.450, and permuting which residue owns which property value, leaving every contact untouched, gives 0.448 ± 0.045 against an observed 0.406 (z = −0.93). Randomized properties perform as well as real ones. The signal was ligand size.

#### Per-residue contact patterns

Recording which position each ligand touches, without summing, gives ρ = 0.628, or 70% of the ceiling. This construction does survive the permutation control: shuffling residue identity within each compound, which preserves that compound’s total contact count exactly, collapses performance to 0.118 ± 0.059 (z = +8.70). The pattern is genuinely positional.

It does not survive size. Residualizing ΔpK_i_ on the five trivial descriptors within each training fold leaves the contact pattern at ρ = 0.037, indistinguishable from its own size-preserving shuffle (−0.002 ± 0.047, z = +0.85), while the Morgan fingerprint retains 0.258 on the same residual. Adding the pattern to the fingerprint changes nothing (−0.015), and under full residualization the combination is worse than the fingerprint alone (0.172 against 0.258).

Position and molecular size are themselves correlated: a larger ligand occupies more of the pocket and therefore contacts a different and larger set of positions. The contact pattern is positional, and its positional content is size.

#### The conclusion is narrow and it is the sharpest statement this work supports

After molecular size is removed, a real chemical signal remains in these data, ligand descriptors capture roughly a third of the attainable ceiling of it, and nothing we derived from the docked poses, neither the scoring function, nor receptor structural properties, nor the contact pattern itself, recovers any of it. The information distinguishing these subtypes is not absent from the benchmark. It is absent from every structural readout we have been able to construct.

### What these results imply a sufficient method must do

The α_2_ selectivity signal is not in static docking scores, which reach 0.071 to 0.188 and are beaten by a model with no receptor. It is not in receptor structural properties or in the pose contact pattern, both of which reduce to molecular size under control. And a static structural statistic cannot establish that the divergence it localizes is subtype-specific rather than crystallographic.

What the benchmark does contain is size, at 72% of the attainable ceiling from five descriptors that require no structure and no model, and a residual chemical signal of ρ = 0.258 that survives size removal and that ligand descriptors capture. Nothing derived from the docked poses reaches that residual.

A sufficient method therefore has a concrete target rather than an open one. It must exceed 0.645 from trivial descriptors, and it must recover some part of the 0.258 of non-size chemical signal that every structural readout tested here misses entirely. Whether a state-resolved readout can do so is not established here, and evaluating one requires all five controls: a measured ceiling, a ligand-only null, a nonselective reference, a same-receptor floor, and a trivial-descriptor baseline.

The two divergent shell positions sit at the rim where the pocket opens, adjacent to the most mobile element of the structure, which is where a state-resolved readout would have to look. That localization is suggestive and, by Calibration 4, not demonstrated.

## Discussion

### An α2-specific result, in a regime where the usual shortcut is unavailable

The same modeling stack that is uninformative on α_2A_/α_2C_ performs respectably on dopamine D_3_/D_2_, and the temptation is to read that as a difference in difficulty. The measurement says something more specific. A model given nothing but Butina cluster membership, no chemistry, no structure, only which series a compound belongs to, explains half the variance in D_3_/D_2_ selectivity and none of the variance in α_2_ selectivity, and calls the D_3_-selective class at AUC 0.852 against 0.489 for α_2_. The two benchmarks are not the same task. On D_3_/D_2_, a substantial part of the target can be reached by recognizing which chemical series a compound belongs to; on α_2_, that route is closed, and 108 compounds occupying 89 clusters leave almost no series structure to exploit.

This matters for how a method’s performance should be read rather than for whether any particular result is correct. A structure-based method scored on D_3_/D_2_ is being asked to do something it can partly avoid doing, and its score will reflect both the structural reasoning it performs and the series recognition it does not need to perform. Reporting a cluster-identity-only baseline alongside any selectivity model separates the two, and it costs one additional fit.

α_2_ is the harder regime, and that is what makes it the informative case. When a ligand-only fingerprint model beats both a physics-based docking score and a CNN rescorer on the same compounds and the same labels, the shortfall cannot be attributed to a benchmark that was solvable without structure.

### Difference endpoints and the independence assumption

The ceiling correction is the result with the widest reach, and its scope should be stated precisely. We did not find that a propagated ceiling is mis-specified in general. We found that the assumption behind it, that the errors in the two affinities being differenced are independent, does not hold for these data, and measured how much difference that makes.

The condition is checkable and, in this benchmark, strongly met: 525 of 586 compounds carry both of their affinities from a single source document, so the two measurements share laboratory, protocol, batch and radioligand, and their errors partly cancel in the difference. Under that structure the propagated variance 2σ^2^ overstates the true noise by roughly a factor of 2.6, and the attainable Spearman ceiling moves from 0.704 to 0.897. The correction replicates on an independently assembled curation with 2.3-fold less evidence, giving 0.520 → 0.819 with a within-document error correlation of 0.633 against 0.659.

Whether other selectivity datasets share this provenance structure is an empirical question we do not answer here. The relevant point is that it is cheap to answer: the estimator in Equation (3) needs only compounds measured in more than one document, requires no assumption about the correlation, and was validated against a Monte Carlo target. A dataset whose paired values come predominantly from single documents is in the correlated-error regime by construction, and the propagated ceiling will understate what is attainable there.

The direction of the error is worth noting because it is not symmetric in its consequences. An under-measured ceiling makes weak predictors look closer to the limit of what is possible than they are. In this benchmark it would have placed the ligand-only null at apparent saturation and left the impression that little headroom remained. Against the measured ceiling, the strictest ligand-only null reaches 62.9%, so a substantial fraction of the attainable signal is being captured by neither chemistry nor static structure.

### What a contact-divergence statistic can and cannot establish

Our own extracellular-enrichment result illustrates a limit that applies to the inference, not to any particular application of it. Subtype-divergent contacts between α_2A_ and α_2C_ are enriched 1.88-fold in the extracellular region, and against a permutation null that randomizes contact placement this is emphatic at p = 0.00005. The detector is not simply manufacturing signal: applied unchanged to D_3_/D_2_, where a secondary binding pocket is independently established, it localizes divergence at 1.60-fold.

The control we had not run is the same analysis on two independent structures of one receptor, where subtype divergence is zero by construction. Those comparisons reach 1.77-fold and 1.68-fold at nominally significant p-values. Against that floor the α_2_ value is not distinguishable: four interpreted cross-subtype pairs average 1.51 against 1.54 for the three same-receptor pairs, and one genuine cross-subtype pair shows no enrichment at all.

The interpretation is mechanical rather than surprising once stated. Extracellular loops are the most conformationally variable region of any two class A GPCR structures, and they differ between two crystals of the same protein for reasons that have nothing to do with subtype: construct design, crystallization conditions, resolution, modeling of weak density. A permutation null over contact placement cannot represent that source of variation, because it holds the structures fixed and randomizes only the assignment. A same-receptor pair can, because it contains exactly that variation and no subtype signal.

We therefore report the α_2_ enrichment as not established rather than as a localization result, and we supply the floor as the control that decides the question. We make no statement about applications of this class of statistic that we did not examine.

Two properties of the comparison set its resolution. The structural comparison rests on one pair of inactive-state structures bound to the same antagonist, which holds ligand identity constant and makes the pair unusually clean, and which confines the conclusion to that state. The floor rests on three same-receptor pairs, so it is the existence of values at 1.68 and 1.77 that carries the argument rather than the mean of the three, and the β_1_/β_2_ comparison is set aside because β_1_ is the turkey receptor and its ECL2 landmark sits mid-axis.

### The target for a sufficient method

The two divergent shell positions sit at the rim where the orthosteric pocket opens into the vestibule, adjacent to ECL2, the most mobile element of the structure and the one a single crystal conformer represents least well. That coincidence is suggestive rather than demonstrated, and it points to where a state-resolved readout would have to look.

The competence controls delimit what the scoring results can be attributed to. Blind self-redocking and the state classifier establish that pose recovery and state assignment are working, which removes setup as an explanation; they measure search competence rather than prospective accuracy, since the engine may have encountered these structures and the box derives from the native ligand’s contacts. Calibration 3 likewise supports a design requirement rather than a free energy: it rests on one run carrying five documented defects, two of them bookkeeping, and Glide XP is no longer available under our license, so those calculations stand as generated in July 2026.

Testing such a readout requires all five controls established here. Without a measured ceiling the comparison is against a misplaced target; without a ligand-only null it is not clear that structure contributes anything; without a nonselective reference an effect-sized number can be manufactured; without a same-receptor floor a structural statistic cannot be shown to reflect subtype rather than crystallography; and without a trivial-descriptor baseline a result that is entirely molecular size will read as structural insight. Whether a state-resolved method clears all five is not established here.

The target is now specific rather than open. A sufficient method must exceed ρ = 0.645 from five descriptors that need no structure, and must recover part of the ρ = 0.258 of non-size chemical signal that every structural readout tested here misses. That is a harder bar than the one implied by comparing against zero, and it is the bar the data supports.

## Conclusions

Subtype selectivity among α_2_-adrenergic receptors is not explained by ligand chemotype, which reaches R^2^ = 0.419, nor by static structure-based scoring, which reaches Spearman 0.071 for Glide SP and 0.188 for CNN rescoring and is outperformed by a ligand-only model that never sees a receptor. The structural reason is visible in the coordinates: 16 of the 18 residues lining the orthosteric pocket are shared between the subtypes, and the two that differ, Glu189→Gly203 and Ile190→Leu204, sit adjacent at the pocket’s extracellular rim rather than in its interior.

The transferable contribution is five controls, each of which changed a conclusion in this work. A directly measured rather than propagated noise ceiling, which requires only compounds measured in more than one document and raises the attainable correlation here from 0.704 to 0.897. A cluster-identity-only null, which quantifies how much of a selectivity benchmark is solvable by series recognition alone and separates D_3_/D_2_ (R^2^ = 0.499) from α_2_ (−0.017). A nonselective reference ligand, without which a free-energy protocol assigned between +1.43 and +4.79 kcal/mol of apparent selectivity to a ligand that has none. A same-receptor technical floor, against which a contact-divergence statistic that appeared highly significant is no longer distinguishable from crystallographic variation. And a trivial-descriptor baseline: molecular weight, heavy-atom count, rotatable bonds, polar surface area and logP together reach 72% of the attainable ceiling, so any method scored on a benchmark of this kind competes first against molecular size.

For α2-directed drug discovery the practical implications are immediate. Programs seeking α_2A_ selectivity (the subtype carrying the withdrawal-suppressing action of clonidine and lofexidine, against the α_2B_-mediated vascular liability) should not expect static docking scores to rank candidates, and should recognize that a ligand-only model will outperform them on retrospective benchmarks without providing any structural rationale to design against. The same caution applies to the α_2_ agonists now prominent as adulterants in the illicit opioid supply, xylazine and medetomidine, where subtype attribution of effects is a live pharmacological question and no crystal structure of either complex exists. Both dock into the two orthosteric sites in the expected pose, anchored by the conserved Asp3.32 salt bridge, in receptors whose pockets clear the native-ligand controls (Figure S5); by the result above, the scores those poses carry are not a basis for assigning subtype preference. Where a selectivity determinant is known and structurally characterized, as in the D_3_ secondary pocket, a static score reproduces its direction and about a third of its magnitude; where it is not, as in α_2_, the score is uninformative.

More generally, the five controls are cheap relative to the calculations they qualify. Measuring a ceiling costs one pass over the provenance of an existing dataset; a cluster-identity baseline costs a single fit; a nonselective reference doubles a free-energy calculation; a same-receptor floor requires one additional structure pair; and a trivial-descriptor baseline costs a few seconds from SMILES, yet in this benchmark it reaches 72% of the attainable ceiling. Each is small next to the cost of a method development program directed at a target that a properly calibrated benchmark would have shown was already saturated, or of a selectivity free energy that a nonselective reference would have shown was manufactured.

## Acknowledgements

This publication was made possible by an Institutional Development Award (IDeA) from the National Institute of General Medical Sciences of the National Institutes of Health under Grant # 2P20GM103432. The content is solely the responsibility of the authors and does not necessarily represent the official views of the National Institutes of Health.

## Author contributions

M Nael and K Elokely were involved in the conception and design of this work. M Nael and K Elokely took part in the analysis and interpretation of the data. M Nael and K Elokely drafted the paper and revised it critically for intellectual content. All authors have given their final approval of the version to be published and agree to be accountable for all aspects of the work.

## Data availability

The frozen evaluation set, the paired dopamine comparison set, and every intermediate artifact behind a reported number are provided in the Supporting Information. The analysis code is available from the corresponding author on request.

