## Supplementary material for "Molecular size dominates α_2_-adrenergic subtype-selectivity benchmarks: five controls for reducing attrition in selective ligand design": SI

**Contents**

**Supporting figures**

S1. Full-length superposition of $\alpha_{2A}$ and $\alpha_{2C}$ S2. Within-document error correlation on both curations
S3. Contact-divergence panel against the same-receptor floor
S4. Interior packing differences between the two receptors
S5. Xylazine and medetomidine in the two orthosteric sites

**Supporting tables**

S1. The orthosteric shell, residue by residue
S2. Contact-divergence panel, all eight pairs
S3. Pose- and state-competence controls
S4. Chemotype separability, full statistics
S5. Ceiling estimation on both curations
S6a. Same-receptor floor for global structural properties S6. Glide XP scores for the R-22 comparison
S7. Molecular size and the size-residualized scores

**Supporting notes**

S1. Validation of the direct noise estimator
S2. Pearson closed form versus Spearman ceiling
S3. The five defects of the free-energy calculation
S4. Defining the orthosteric shell
S5. A pre-registered chemotype-stratified test

**Supporting data files**, contents listed at the end.

### Supporting figures

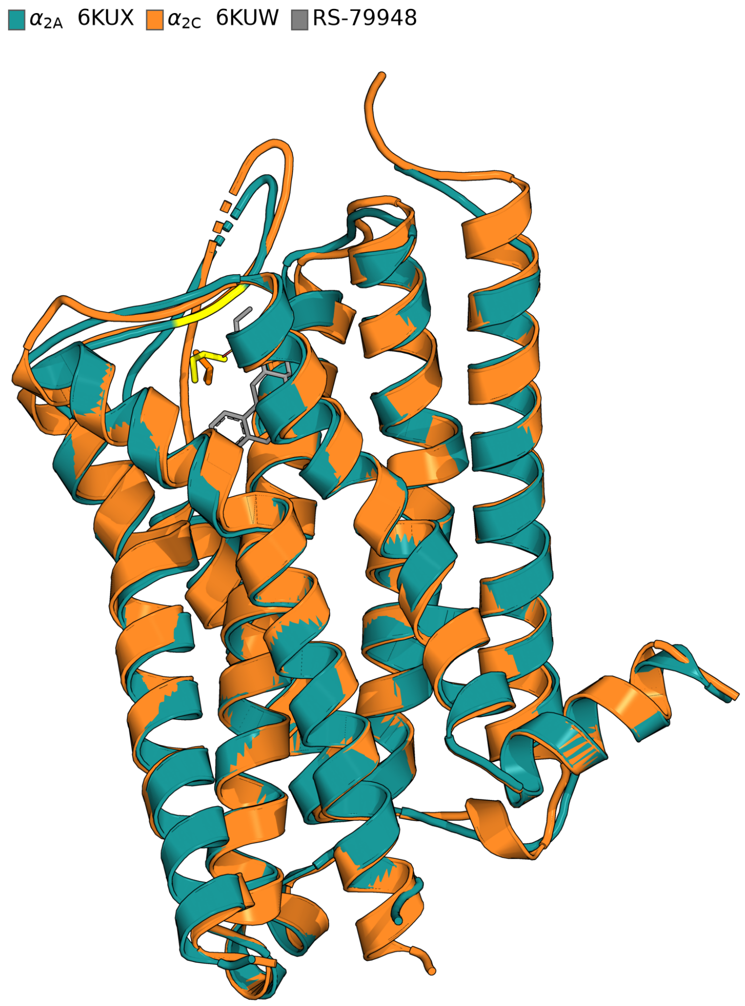


**Figure S1. Full-length superposition of $\alpha_{2A}$ (6KUX, teal) and $\alpha_{2C}$ (6KUW, orange), both bound to RS-79948.** Receptors are shown in the membrane frame, extracellular face up. Gray sticks, the co-crystallized antagonist in $\alpha_{2A}$; yellow and orange sticks, the two divergent orthosteric-shell positions. Sequence-independent superposition (super) gives 0.743 Å over 1,209 atom pairs; sequence-aware superposition with outlier rejection (align) gives 0.638 Å over 1,425 pairs, from 2.093 Å over 1,907 pairs before rejection. The figure is included to document the extent of global similarity; the informative comparison is local and appears as Figure 1.

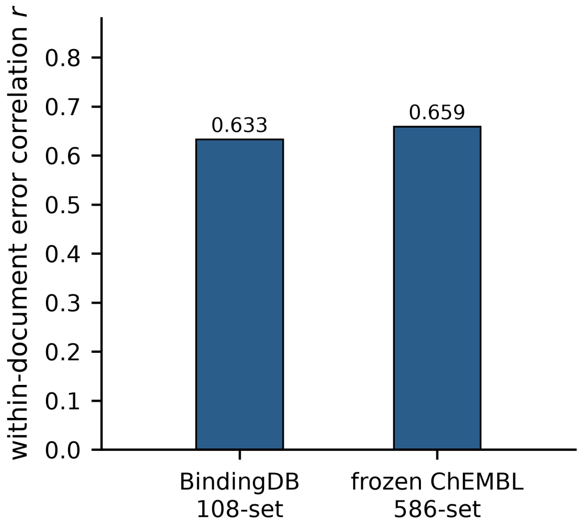


**Figure S2. Within-document error correlation.** The correlation between $\alpha_{2A}$ and $\alpha_{2C}$ measurement errors for compounds whose two affinities are reported in the same source document, measured independently on the two curations: r = 0.633 on the BindingDB 108-set and r = 0.659 on the frozen ChEMBL 586-set. Under independence this quantity would be zero, and the propagated noise variance $2\sigma^2$ would be correct.

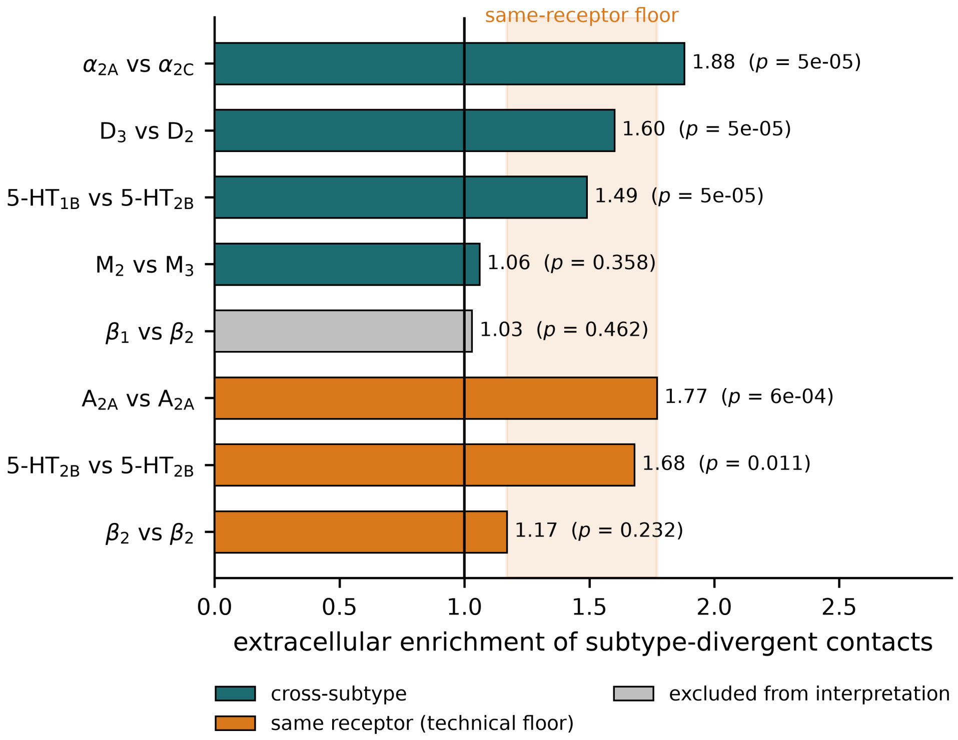


**Figure S3. A contact-divergence statistic does not clear its own same-receptor floor.** Extracellular enrichment of subtype-divergent contacts for eight construct-harmonized receptor pairs, with permutation p-values. Teal, cross-subtype comparisons; orange, comparisons between two independent structures of the same receptor, where subtype divergence is zero by construction; grey, the $\beta_1/\beta_2$ pair, excluded from interpretation because $\beta_1$ is the turkey receptor and its ECL2 landmark sits mid-axis. The shaded band spans the same-receptor values and marks the technical floor. The vertical line at 1.0 is the enrichment expected if divergent contacts were distributed as the background contacts are.

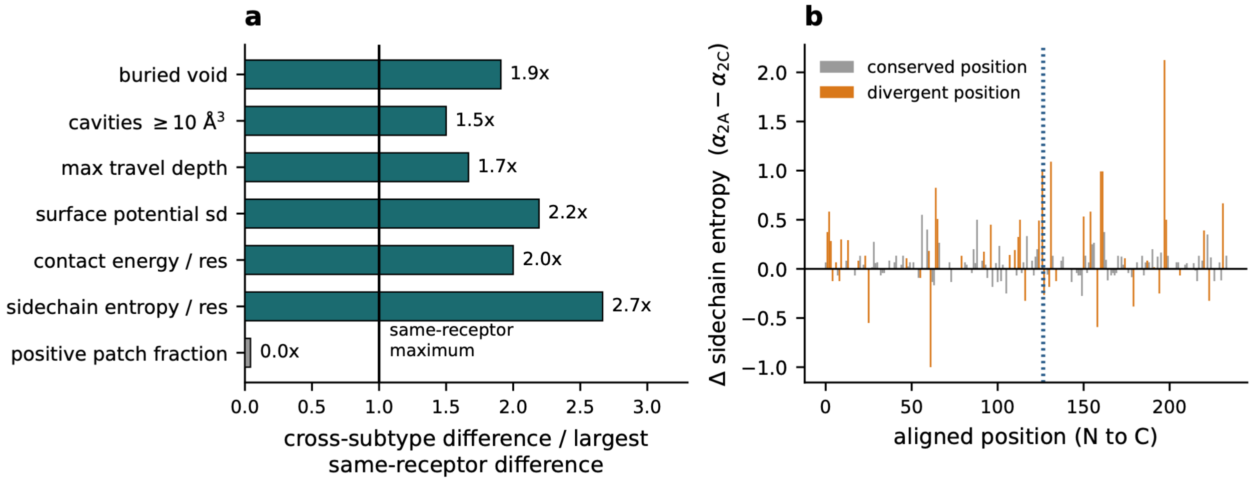


**Figure S4. The two receptors differ in interior packing by margins that three same-receptor pairs do not reproduce.** (a) Each cross-subtype property difference divided by the largest difference seen among three pairs of independent structures of a single receptor ($\beta_2$ 2RH1/3NY8, 5-$HT_{2B}$ 4IB4/4NC3, $A_{2A}$ 3EML/4EIY), all harmonized identically. Teal, above the same-receptor maximum; grey, at or below it. The vertical line marks that maximum. The floor comprises three pairs and the properties are correlated, so the margins are read as suggestive; the full table is Table S6a. (b) Difference in per-residue sidechain entropy along the sequence alignment, $\alpha_{2A}$ minus $\alpha_{2C}$. Orange, positions where the two receptors carry different residues; grey, positions where they are identical. The dotted line marks the two divergent orthosteric-shell positions, Glu189/Gly203 and Ile190/Leu204.

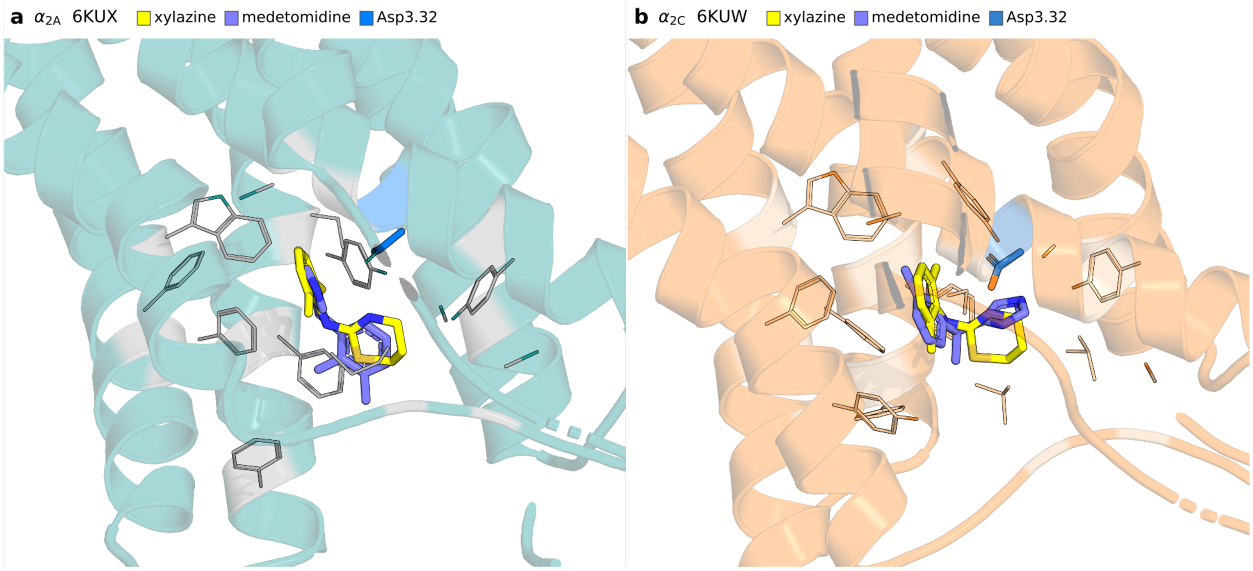


**Figure S5. Xylazine and medetomidine in the $\alpha_{2A}$ and $\alpha_{2C}$ orthosteric sites.** (a) $\alpha_{2A}$ (6KUX, teal) and (b) $\alpha_{2C}$ (6KUW, orange), each with xylazine (yellow) and medetomidine (slate) docked as the protonated amidinium. Shell residues within 5.0 Å of either ligand are shown as thin lines; the conserved Asp3.32 anchor is blue in both panels. Non-polar hydrogens are hidden. Both receptors are the crystal structures whose pockets clear the native-ligand controls (score-only -7.75 and -8.82 kcal/mol; native re-docking to 0.93 and 0.80 Å). Poses are the best-scoring ones meeting the Asp3.32 salt-bridge and clash criteria: xylazine -6.76 and -6.81 kcal/mol with salt bridges of 2.96 and 3.08 Å, medetomidine -7.12 and -8.15 kcal/mol with 3.29 and 3.25 Å. Given that a docking score on this benchmark is outperformed by molecular weight, these values are shown as poses rather than as predicted selectivities.

### Supporting tables

#### Table S1. The orthosteric shell, residue by residue

All 18 residues with at least one heavy atom within 5.0 Å of the co-crystallized RS-79948 in 6KUX, mapped through the sequence alignment to their $\alpha_{2C}$ counterparts. Machine-readable version: TableS1_orthosteric_shell.csv.

| $\alpha_{2A}$ | $\alpha_{2C}$ | status |
|---|---|---|
| Ser90 | Ser108 | conserved |
| Tyr109 | Tyr127 | conserved |
| **Asp113** (3.32) | **Asp131** | conserved |
| Val114 | Val132 | conserved |
| Cys117 | Cys135 | conserved |
| Thr118 | Thr136 | conserved |
| Cys188 (ECL2) | Cys202 | conserved |
| **Glu189** | **Gly203** | **divergent** |
| **Ile190** | **Leu204** | **divergent** |
| Ser200 (5.42) | Ser214 | conserved |
| Ser204 (5.46) | Ser218 | conserved |
| Trp387 (6.48) | Trp395 | conserved |
| Phe390 (6.51) | Phe398 | conserved |
| Phe391 (6.52) | Phe399 | conserved |
| Tyr394 (6.55) | Tyr402 | conserved |
| Phe408 | Phe419 | conserved |
| Phe412 | Phe423 | conserved |
| Tyr416 (7.43) | Tyr427 | conserved |

The canonical aminergic recognition machinery is conserved in its entirety: the Asp3.32 salt-bridge anchor, both TM5 serines, the Trp6.48 toggle, the Phe6.51/Phe6.52 aromatic cage, Tyr6.55 and Tyr7.43. Cys188 and Cys202 are the ECL2 partners of the conserved Cys3.25-CysECL2 disulfide. The two divergent positions are adjacent, immediately following that cysteine, at the rim where the pocket opens into the vestibule.

**Table S2. Contact-divergence panel, all eight pairs**

Machine-readable version: contact_divergence_panel_8pairs.json.

| pair | kind | changed contacts | background | in region (%) | background (%) | enrichment | permutation p |
|---|---|---|---|---|---|---|---|
| $\alpha_{2A}$ vs $\alpha_{2C}$ | cross-subtype | 132 | 1,229 | 52.3 | 27.7 | 1.88 | 0.00005 |
| $D_3$ vs $D_2$ | cross-subtype | 151 | 1,270 | 55.6 | 34.7 | 1.60 | 0.00005 |
| 5-$HT_{1B}$ vs 5-$HT_{2B}$ | cross-subtype | 250 | 1,242 | 40.4 | 27.1 | 1.49 | 0.00005 |
| $M_2$ vs $M_3$ | cross-subtype | 92 | 1,290 | 41.3 | 39.1 | 1.06 | 0.358 |
| $\beta_1$ vs $\beta_2$ | cross-subtype, excluded | 98 | 1,275 | 28.6 | 27.8 | 1.03 | 0.462 |
| **$A_{2A}$ vs $A_{2A}$** | **same receptor** | 64 | 1,357 | n/a | n/a | **1.77** | 0.00055 |
| **5-$HT_{2B}$ vs 5-$HT_{2B}$** | **same receptor** | 45 | 1,312 | n/a | n/a | **1.68** | 0.011 |
| $\beta_2$ vs $\beta_2$ | same receptor | 60 | 1,335 | n/a | n/a | 1.17 | 0.232 |

Mean over the four interpreted cross-subtype pairs, 1.51; mean over the three same-receptor pairs, 1.54. $\beta_1/\beta_2$ is excluded from interpretation: $\beta_1$ is the turkey receptor and its ECL2 landmark sits mid-axis, making the geometric region assignment unreliable. It is retained in the figure, grayed, so that the full panel is visible.

**Table S3. Pose- and state-competence controls**

**Gate 1, blind self-redocking**, five $\alpha_2$ co-crystal complexes, box from the contact frame rather than the native ligand, hardened to 20 seeds. Metric is symmetry-corrected heavy-atom RMSD of the top-ranked pose.

| PDB | subtype | state | ligand | median top-1 RMSD (Å) | seeds ≤ 2 Å |
|---|---|---|---|---|---|
| 7W7E | $\alpha_{2A}$ | active | biased agonist | 0.37 | 20/20 |
| 9CBL | $\alpha_{2A}$ | active | epinephrine | 0.25 | 20/20 |
| 6KUX | $\alpha_{2A}$ | inactive | RS-79948 | 0.63 | 20/20 |
| 6K41 | $\alpha_{2B}$ | active | dexmedetomidine | 0.42 | 20/20 |
| 6KUW | $\alpha_{2C}$ | inactive | RS-79948 epimer | 0.73 | 20/20 |

**Gate 2, structure-only state assignment.** Score is the TM3-TM6 Cα distance between positions 3.50 and 6.34; the 10 Å threshold was fixed in advance from the literature and not tuned on these structures. 7W7E and 6KUX were used as references and the remaining three assigned blind.

| PDB | truth | TM3-TM6 (Å) | call | blind |
|---|---|---|---|---|
| 7W7E | active | 16.08 | active | reference |
| 9CBL | active | ~16 | active | yes |
| 6K41 | active | 15.92 | active | yes |
| 6KUX | inactive | 6.83 | inactive | reference |
| 6KUW | inactive | 6.31 | inactive | yes |

Blind calls 3/3; 5/5 overall. Both gates measure search and setup competence only, not prospective accuracy: the docking engine may have been exposed to these structures during development, and the search box derives from the native ligand's own contacts.

**Table S4. Chemotype separability, full statistics**

Machine-readable version: chemotype_separability.json. Butina clustering of Morgan fingerprints, Tanimoto cutoff 0.35, applied identically to both pairs.

| | $\alpha_{2A}/\alpha_{2C}$ | $D_3/D_2$ |
|---|---|---|
| compounds / clusters | 108 / 89 | 934 / 226 |
| compounds per cluster | 1.2 | 4.1 |
| **$T_1$ cluster-identity-only $R^2$** | **−0.017** | **+0.499** |
| $T_2$ ECFP random-split CV $R^2$ | +0.437 | +0.705 |
| $T_2$ ECFP cluster-grouped CV $R^2$ | +0.347 | +0.639 |
| $T_2$ Δ (within-series memorization) | +0.090 | +0.066 |
| **$T_3$ cluster-only class AUC** | **0.489** | **0.852** |
| $\eta^2$ between-cluster variance | 0.936 | 0.773 |

$T_2$ does not discriminate between the two pairs and is reported as a null. $\eta^2$ is uninformative at $\alpha_2$'s granularity, where 108 compounds occupy 89 clusters and between-cluster variance is near-total by construction; it is reported for completeness and is not used as a test. $T_1$ and $T_3$ are cross-validated and immune to that artifact. $D_3/D_2$ composition: 575 $D_3$-preferring, 5 $D_2$-preferring, 354 nonselective.

**Table S5. Ceiling estimation on both curations**

Machine-readable versions: ceiling_estimation_bindingdb108.json, frozen_set_rescore.json.

| | BindingDB 108-set | frozen ChEMBL 586-set |
|---|---|---|
| compounds with a same-document matched pair | 91/106 (86%) | 525/586 (90%) |
| compounds with ≥ 2 paired documents | 24 | 55 |
| cross-document pair-differences | 39 | 136 |
| within-document error correlation r | 0.633 | 0.659 |
| measured noise variance (Eq. 3) | 0.165 | 0.139 |
| propagated noise variance $2\sigma^2$ | 0.366 | 0.360 ($\sigma$ = 0.424) |
| **$\rho_{max}$ measured** | **0.819** [0.715, 0.944] | **0.897** [0.844, 0.960] |
| $\rho_{max}$ propagated | 0.520 | 0.704 |

The measured noise is 45% and 39% of the propagated value respectively. Confidence intervals are bootstrap over compound clusters (n = 24 clusters, ≈ 30 effective degrees of freedom on the BindingDB set), not over the individual (compound, document) deviations, which are not independent. The upper bound of 0.944 should not be quoted as a candidate ceiling; it implies r = 0.85.

**Table S6a. Same-receptor floor for global structural properties**

Absolute difference within each pair, all structures harmonized identically. Machine-readable version: property_floor.json.

| property | $\alpha_{2A}/\alpha_{2C}$ | $\beta_2$ (2RH1/3NY8) | 5-$HT_{2B}$ (4IB4/4NC3) | $A_{2A}$ (3EML/4EIY) |
|---|---|---|---|---|
| buried void (Å^3) | 166 | 33 | 25 | 87 |
| cavities >= 10 Å^3 | 3 | 0 | 1 | 2 |
| maximum travel depth (Å) | 7.5 | 4.5 | 3.0 | 1.5 |
| surface potential sd | 134.6 | 5.5 | 2.4 | 61.4 |
| contact energy per residue | 0.34 | 0.14 | 0.04 | 0.17 |
| sidechain entropy per residue | 0.08 | 0.01 | 0.00 | 0.03 |
| positive patch fraction | 0.01 | 0.10 | 0.24 | 0.01 |
| modeled residues | 269/278 | 282/279 | 285/287 | 288/299 |

Six of seven cross-subtype differences exceed the largest same-receptor value. The floor is three pairs, the properties are correlated, and the $A_{2A}$ pair differs in modeled length; see the main text.

**Table S6. Glide XP scores for the R-22 comparison**

Best-scoring pose per ligand per receptor. Raw output including all XP descriptor terms and both protomers of each ligand: glide_xp_R22_eticlopride_raw.csv.

| ligand | $D_3$ (kcal/mol) | $D_2$ (kcal/mol) | difference |
|---|---|---|---|
| R-22 | −9.580 | −7.971 | −1.609 |
| eticlopride (nonselective) | −6.969 | −6.551 | −0.418 |
| **baseline-corrected** | | | **−1.190** |
| experimental (566-fold, 310 K) | | | **−3.906** |

R-22 was supplied in two protonation states; the tabulated value is the better-scoring pose in each receptor. No percentage recovery is reported, because equating a GlideScore difference with a binding free energy is the assumption under examination.

**Table S7. Molecular size and the size-residualized scores**

Machine-readable versions: join_results.json, size_control.json, pattern_results.json, mw_residual.json. All Spearman correlations against measured dpKi on the frozen set under scaffold-grouped cross-validation, except the raw descriptor correlations, which need no fitting.

**What size alone achieves**

| predictor | rho |
|---|---|
| molecular weight (no model) | -0.561 |
| heavy-atom count (no model) | -0.575 |
| ligand heavy atoms in contact, $\alpha_{2A}$ (no model) | -0.505 |
| MW only, cross-validated | 0.564 |
| MW + heavy atoms | 0.608 |
| MW + heavy atoms + rotatable bonds + TPSA + logP | **0.645** |

**Feature sets and their controls**

| feature set | rho | control | control rho |
|---|---|---|---|
| contact-weighted receptor properties | 0.406 | size only, six contact counts | 0.450 |
| | | properties permuted across residues | 0.448 +/- 0.045 |
| | | size-normalized construction | 0.400 |
| per-residue contact pattern, both receptors | 0.628 | residue identity shuffled within compound | 0.118 +/- 0.059 |
| ligand fingerprint | 0.741 | | |

**After molecular size is removed** (size model fitted within each training fold)

| feature set | MW only | MW + heavy | full five-descriptor block |
|---|---|---|---|
| contact pattern | +0.152 | +0.114 | **+0.037** |
| ligand fingerprint | +0.293 | +0.268 | **+0.258** |
| pattern + fingerprint | +0.275 | +0.240 | +0.172 |

On the strictest residual the column-shuffled contact pattern gives -0.002 +/- 0.047 against an observed +0.037. The pattern is positional before size is removed and is not separable from its size-preserving shuffle afterwards, while the fingerprint retains 0.258.

### Supporting notes

#### Note S1. Validation of the direct noise estimator

The estimator $\sigma^2_{measured} = Var(\Delta_i - \Delta_j)/2$ makes no assumption about the correlation between the two affinity errors, but it does assume the two documents contribute independent errors. It was validated by Monte Carlo: simulating from the empirical label distribution with a known injected noise variance of 0.16503, the estimator recovered 0.16532 over 4,000 replicates.

Three challenges to the estimate were tested directly. First, that the compounds carrying the estimate might be cleaner than the benchmark as a whole: within-subtype σ across the 24 estimating compounds is 0.4173, against 0.4258 for all replicated compounds and 0.4275 globally, a ratio of 0.976, so they are not measurably cleaner. Second, that median aggregation across multiple documents could alone explain the discrepancy: even after a generous median-aware correction, the independence-based ceiling reaches only 0.563, still below the 0.605 ligand-only null on that set, so the correlated-error correction remains load-bearing. Third, the Pearson-versus-Spearman question, treated in Note S2.

We also record what the agreement between the direct estimate (0.819) and an earlier split-half estimate (0.817) is not. Both are computed on essentially the same 24 to 25 best-replicated compounds and the same between-document scatter, so their agreement is an internal consistency check on the algebra, not independent corroboration. The independent evidence is the replication on the frozen ChEMBL set.

#### Note S2. Pearson closed form versus Spearman ceiling

Equation (4) is a Pearson attenuation result, while every score reported in the manuscript is a Spearman rank correlation. The two were reconciled by simulation on the empirical $\Delta pK_i$ distribution, with noise scaled per compound by the number of contributing documents (k-distribution: 61 singletons, 18 with $k = 2$, 11 with $k = 3$, 16 with $k \geq 4$; mean $1/k = 0.724$).

| noise model | Pearson closed form | Spearman, single-document noise | Spearman, median-aware |
|---|---|---|---|
| same-document (0.1650) | 0.819 | 0.791 | 0.837 |
| blended, conservative (0.1934) | 0.783 | 0.755 | 0.806 |
| original independence (0.3655) | 0.520 | 0.493 | 0.563 |

The empirical label distribution is close to normal, so the heavy-tail penalty is 0.01 to 0.02 rather than the 0.10 to 0.15 that might be feared, and the median-aggregation correction of about +0.045 acts in the opposite direction and is slightly larger. The values reported in the manuscript are therefore mildly conservative.

**Note S3. The five defects of the free-energy calculation**

Calibration 3 is reported as a design requirement rather than as a measurement. The defects, in full:

1. **Not robust to analysis choice.** Across nine definitions formed by crossing three bulk references (1.8, 1.9, 2.0 nm) with three orthosteric boundaries (0.8, 0.9, 1.0 nm), the nonselective control receives +1.43 to +4.79 kcal/mol, a spread of 3.36 kcal/mol, as large as the effect modeled.
2. **Not converged.** The free-energy surface was still filling at 150 ns per walker. Over the final 20%, the span grew by about +1.6 kcal/mol in the $D_3$ eticlopride system against +0.91 in $D_2$, leaving roughly 0.7 kcal/mol of differential still accumulating.
3. **The conventional convergence diagnostic is vacuous here.** Multiple-walker dispersion is 0.004 to 0.058 kcal/mol, but the three walkers share a single bias grid, so their agreement is near-perfect by construction and reports sub-0.1 kcal/mol precision on a number that moves by 3.4 kcal/mol under analysis choice.
4. **No time-block test is possible.** The surface file was overwritten every 100 ps; only the final surface was retained.
5. **The topologies carry no disulfide bonds.** In all three $D_3$ systems, two cysteine pairs sit at 2.03 Å in the coordinates (Cys72-Cys150, the conserved Cys3.25-CysECL2 bond, and Cys227-Cys230), yet neither is bonded in the force field, leaving 13 free thiols per system. That bond tethers ECL2 to TM3 at the extracellular vestibule the reaction coordinate traverses.

A sixth point limits the supporting observation rather than the control: SB-277011A had no crystal pose, and its geometry is a docked conformer, so its near-zero value indicates inversion but cannot carry that claim alone.

**Note S4. Defining the orthosteric shell**

S5. A pre-registered chemotype-stratified test

The shell must be selected as whole residues within the distance cutoff. Selecting atoms within 5.0 Å and then filtering to Cα retains only those residues whose Cα itself falls inside the cutoff and returns 3 residues rather than 18, a pocket that excludes Asp3.32 and every aromatic contact. The reported shell uses byres expansion of the 5.0 Å atom shell.

**Note S5. A pre-registered chemotype-stratified test**

The pooled analysis reports that ligand chemotype explains part of $\alpha_2$ selectivity and that static docking explains none of it beyond a ligand-only null. Both figures are pooled over a chemically heterogeneous set, and pooling would conceal a determinant operating within one chemotype. We tested this with the criteria fixed in advance.

**Registration.** The protocol was written and hashed (SHA-256 8693c83960e56fed3243500abded3aa2ef5a212d7ae602d859775809f7b783fc) before any stratified quantity was computed; it is included as PREREG_STAGE1.md. Strata were defined by SMARTS from the medicinal chemistry of $\alpha_2$ ligands, deliberately not from the Butina clusters used elsewhere, since clusters derived from the same fingerprints the null model uses would make strata and null circular. Strata entered the analysis at n >= 30. The registered criterion was a permutation test on the stratum labels, requiring **both** p_strat < 0.01 **and** a compound-level cluster bootstrap confidence interval on the winning stratum excluding zero, with a stopping rule specifying that Stages 2 and 3 would not run otherwise.

**Result.**

| stratum | n | rho Glide SP | rho ligand-only null | delta |
|---|---|---|---|---|
| arylpiperazine | 68 | +0.149 | +0.770 | -0.621 |
| imidazole | 48 | -0.206 | +0.622 | -0.827 |
| imidazoline | 143 | +0.248 | +0.045 | **+0.203** |

Delta_max = +0.203 in the imidazoline stratum. Over 2,000 label permutations preserving stratum sizes, p_strat = 0.0030 (permuted Delta_max mean -0.372, 95th percentile -0.114). The cluster bootstrap on that stratum gave delta = +0.203 with a 95% interval of [-0.179, +0.617].

**One criterion of the two was met, so by the registered stopping rule Stages 2 and 3 were not run.** The permutation asks whether the chemotype partition is special and answers yes; the bootstrap asks whether the effect within imidazolines is estimable and answers that it is not, since 143 compounds in few scaffold groups admit values from -0.179 to +0.617. Only the second licenses a claim about imidazolines.

Two observations recorded afterwards bear on the interpretation. Within the imidazoline stratum the Glide score correlates with molecular weight at -0.323 and molecular weight with dpKi at -0.352, an indirect path accounting for roughly half of the observed delta; the registration predates the size analysis and contains no control for it. And the delta is carried by the null descending rather than by docking ascending: the ligand-only null reaches only 0.045 there, against 0.248 for Glide, which is 28% of the measured ceiling.

The descriptive breakdown is the more useful output, and nothing is gated on it:

| stratum | n | var(dpKi) | ligand-only rho |
|---|---|---|---|
| arylpiperazine | 68 | 0.874 | 0.770 |
| imidazole | 48 | 0.345 | 0.622 |
| other | 311 | 0.480 | 0.520 |
| imidazoline | 143 | 0.252 | 0.045 |

Imidazolines carry both the lowest selectivity variance and a ligand-only null at chance. A tight congeneric series with little selectivity spread gives a fingerprint model little to discriminate on, which accounts for the apparent delta without requiring the docking score to be informative.

**Supporting data files**

| file | contents |
| --- | --- |
| TableS_frozen_set_586_compounds.csv | the frozen evaluation set: compound identifier, standardized SMILES, $pK_i$ at both subtypes, $\Delta pK_i$, scaffold, and per-receptor docking scores (586 rows) |
| TableS1_orthosteric_shell.csv | the 18 shell residues with $\alpha_{2A}/\alpha_{2C}$ correspondence and conserved/divergent status |
| d3d2_paired_934_compounds.csv | the paired dopamine $D_3/D_2$ comparison set |
| ceiling_estimation_bindingdb108.json | all quantities entering the ceiling estimate on the BindingDB curation |
| frozen_set_rescore.json | every method score and both ceilings on the frozen set |
| chemotype_separability.json | T1, T2, $T_3$ and $\eta^2$ for both receptor pairs |
| contact_divergence_panel_8pairs.json | changed contacts, background, enrichment and permutation p for all eight pairs |
| contact_divergence_pairs.csv | the structure pairs and PDB identifiers used in the panel |
| structural_comparison_a2A_a2C.json | superposition statistics, the full shell table, and the axial distribution of divergent residues |
| glide_xp_R22_eticlopride_raw.csv | unmodified Glide XP output for both ligands against both receptors, all descriptor terms retained |
| NUMBER_AUDIT.md | provenance record tracing every figure in the manuscript to its primary artifact, including quantities examined and excluded |
| property_floor.json | same-receptor floor for the global structural properties, all four pairs |
| join_results.json | contact-weighted receptor-property descriptors and their scores |
| size_control.json | size-only baseline, property-shuffle null and size-normalized descriptors |
| pattern_results.json | per-residue contact-pattern features and the column-shuffle control |
| mw_residual.json | size-residualized scores for every feature set and size definition |
| PREREG_STAGE1.md | the pre-registration, as hashed before execution |
| stage1_results.json | stratum sizes, both gates, and the descriptive breakdown |
| size_table_S7.json | every row of Table S7 recomputed under one documented protocol by an independent |

| file | contents |
| --- | --- |
| | reimplementation, agreeing with the table to within 0.006 |
| null_seed_spread.json | the ligand-only null refitted across ten fold assignments |
| null_scaffold_ci.json | the scaffold-grouped ligand-only null with a bootstrap interval over scaffold groups |
| adulterant_docking.json | scores, accepted pose indices and Asp3.32 distances for the two adulterants, measured in the frame Figure S5 renders |
